# Impaired transitioning of the FXR ligand binding domain to an active state underlies an FXR-associated PFIC phenotype

**DOI:** 10.1101/2024.02.08.579530

**Authors:** Annika Behrendt, Alex Bastianelli, Jan Stindt, Eva-Doreen Pfister, Malte Sgodda, Tobias Cantz, Sebastian Hook, Kathrin Grau, Stefanie Brands, Carola Dröge, Amelie Stalke, Michele Bonus, Sabine Franke, Ulrich Baumann, Verena Keitel, Holger Gohlke

## Abstract

Nuclear receptor farnesoid X receptor (FXR) acts as a key regulator of bile acid pool homeostasis and metabolism. Within the enterohepatic circulation, reabsorbed bile acids act as FXR agonists, which transcriptionally controls the synthesis and transport of bile acids. Binding occurs in the ligand binding domain (LBD), favoring a conformational change to the active state in which helix 12 interacts with the LBD to form an interaction surface for nuclear co-activators. The homozygous missense variant T296I, identified in a PFIC patient, is located close to the critical helix 12 interaction. Here, we identified reduced transcriptional activity of the variant protein on the downstream targets bile salt export pump (BSEP) and small heterodimer partner (SHP) *in vitro,* within the patient’s liver, and in iPSC-derived hepatic organoids. BSEP-dependent Tauro-DBD transport was impaired in T296I patient-derived organoids, but could be rescued via lipid nanoparticle-mediated FXR WT mRNA delivery, indicating the variant is responsible for the identified reduced BSEP expression. Using molecular dynamics simulations, we observed a reduced transitioning from the inactive to the active state for the T296I variant, indicating a molecular mechanism underlying the reduced activity. To our knowledge, this is the first study to describe the conformational change from an inactive to an active state of the FXR LBD. This might be useful for new therapeutic approaches targeting the activation of FXR.

## Introduction

Progressive familial intrahepatic cholestasis (PFIC) is a rare group of genetic disorders that affect the liver’s ability to excrete bile constituents, resulting in impaired bile flow, subsequent intrahepatic cholestasis, and progressive liver damage and failure^2,3^.

Farnesoid X receptor (FXR), encoded by the *NR1H4* gene, is a nuclear receptor (NR) responsive to bile acids (BA) and a key regulator of BA metabolism, playing a pivotal role in maintaining BA homeostasis by controlling BA synthesis, transport, and detoxification^4,5^. Amongst other targets, the *ABCB11* gene (encoding for the bile salt export pump (BSEP)) and the *NR0B2* gene (encoding for the small heterodimer partner (SHP)) are transcriptionally regulated by FXR^4,5^. *NR1H4* variants associated with PFIC (FXR-deficiency, also named PFIC subtype 5) were characterized by a secondary BSEP deficiency, coagulopathy and a rapid progression towards end-stage liver disease^6–9^. While most patients carried bi-allelic protein-truncating variants^6–8,10^, only two *NR1H4*-associated PFIC patients carrying homozygous missense variants have been described to date^8,11^. One patient died on the transplant waiting list due to end-stage liver disease at the age of 9 months (c.557G>A)^8^ while the other patient was successfully transplanted at the age of 8 months (c.887C>T, p.(Thr296Ile), referred to as T296I in the following) and is currently 10 years old^11^. For further information on the patient and clinical data, please refer to the initial report (case 4)^11^. Staining of FXR and the bile salt export pump (BSEP) was found negative in the liver tissue of PFIC patients with protein-truncating *NR1H4* variants^6,10^. To determine the contribution of the homozygous *NR1H4* T296I missense variant to the PFIC phenotype of our patient, we studied the localization and transcriptional activity of the mutated protein *in vitro*. We further used the known intrahepatic cholestasis of pregnancy (ICP)-associated M173T variant^12^ to compare variant effects (Fig. S2 and S3). M173T showed decreased transcription of the downstream FXR targets BSEP and ileal BA binding protein (IBABP) in luciferase assays^12^. To unravel the impact of the T296I variant on the structural dynamics of the protein at an atomic level, we further used molecular dynamics (MD) simulations. MD simulations have proven useful in elucidating the functional mechanisms of protein activity^13^. In particular, nuclear receptors (NRs) have benefited from this in-depth analysis as their functions are often diverse, and subtle changes in ligands can lead to altered conformations and, thus, protein activity^14–16^. The positioning of helix 12 (H12), forming part of the activation function 2 (AF2) surface, is pivotal for NR activity via the recruitment of coregulatory proteins. Coactivators interact with the AF2 surface using a conserved LXXLL motif^17^, while antagonist-bound NRs favor corepressor binding to the AF2 surface with a larger hydrophobic motif and blocking the active positioning of H12^18^. Several MD studies of the LBD of FXR have underlined the importance of H12 positioning^19–21^. However, the transitioning from the inactive to the active conformation as well as the effect of single-site missense variants on the function of FXR have so far not been analyzed by MD studies. Thus, we employed MD simulations to analyze the conformational change from an inactive to the active state and evaluated the impact of the T296I variant both with a localized distance measurement and with regard to its influence on H12 positioning. To our knowledge, this is the first simulation study to explicitly investigate the transitioning between the inactive and active state of the FXR LBD, uncovering the conformational change of H12.

## Materials and Methods

### Plasmids, cloning and mutagenesis

The BSEP promoter plasmid based on pGL3-basic (BSEP^prom^-Luc) was a kind gift from Roche. The human small heterodimer partner (*SHP)* promoter (bases -572 to +10, GenBank Accession Number AF044316)^22^ was amplified by PCR from a healthy human liver genomic DNA pool. DNA sequencing was performed for all cDNAs used (Eurofins). Note that the numbering of the protein variant (T296I) is based on the alpha1 isoform (Uniprot acc. Q96RI1-1). For details on the cloning strategies and plasmids see SI.

### Immunofluorescence staining of HEK293 cells

HEK293 cells seeded onto glass coverslips in 12-well plates were transiently transfected with 1µg each of FXR and retinoic X receptor (RXR) α expression constructs for 48h. After 24h, cells were stimulated with obeticholic acid (OCA, INT-747, 10μM) and 9-cis-retinoic acid (9-cis-RA, 1μM). Cells on coverslips were washed with PBS before fixation with ice-cold methanol (30sec). After blocking in UltraVision protein block (ThermoFisher Scientific) for 30min, cells were stained for 1h at 1:100 with rabbit anti-FXR (H-130; sc-13063, Santa Cruz Biotechnology) followed by staining at 1:250 with goat anti-rabbit-IgG-FITC (Jackson ImmunoResearch) and DAPI at 1:20.000. Coverslips were mounted on microscopic slides using Dako fluorescence mounting medium.

### Western Blot

HEK293 cells seeded into 6-well plates were transiently transfected for 48h with 2µg per well of either WT or mutant *pno*Cherry-FXR as described above and in SI. Membranes were blocked with 5% BSA in TBS-T for 1h before overnight incubation with rabbit anti-FXR (H-130; 1:2.000) and mouse anti-β-actin (ab6276, Abcam; 1:10.000) followed by incubation with goat anti-rabbit-IgG-AlexaFluor 647 and goat anti-mouse-IgG-AlexaFluor 488 (both at 1:5.000). Fluorescent signals were detected using a ChemiDoc MP imaging system (Biorad).

### Luciferase Assay

Luciferase assays were performed using the Dual Luciferase reporter assay (Promega) according to the manufactureŕs instructions. Briefly, HEK293 cells kept in DMEM containing 10% fetal calf serum (FCS) were seeded in 96-well plates at 7.000 cells per well and transfected the next morning with 100ng of BSEP-Luc or SHP-Luc plasmid and 10ng each of FXR and RXR expression plasmids using Fugene HD (Promega) at a 2.5:1 (reagent:DNA) ratio. Where appropriate, plasmids were replaced with equal amounts of their respective empty backbones as a control, and each well also received 5ng of pRL-TK as an internal assay control. Cells transfected with FXRα1/2 expression plasmid were stimulated with CDCA (0, 1, 10, or 25µM), cells transfected with RXRα were stimulated with 1μM 9-cis-retinoic acid (9-cis-RA), cells transfected with both FXRα1/2 and RXRα expression plasmids were stimulated with both ligands.

### Patient iPSC-derived hepatic organoids

Induced pluripotent stem cells (iPSCs) were generated from patient peripheral blood mononuclear cells (PBMCs) with a commercial Sendai virus-based reprogramming system (Cytotune 2.0, ThermoFisher Scientific) and maintained as pluripotent stem cells on CF1 murine feeder cells according to standard conditions. Generation of hepatic organoids was performed as described by Shinozawa et al.^23^ with minor changes. In brief, cells were seeded as single cells on GelTrex coated dishes (TPP) in E8 media containing no FGF2 but 10µM RHO-Kinase Inhibitor (Tocris) with a density of 120.000 cells/cm². After 24h cells were washed and differentiated towards definitive endoderm and subsequently embedded into Matrigel (Corning) domes as described to form iPSC-derived hepatic organoids. After 9-11 days, hepatic organoids were matured and used for subsequent qRT-PCR and BSEP transport analyses. As BSEP-specific substrate, we chose the red-fluorescent Tauro-nor-THCA-24-DBD (Tauro-DBD) substrate which was added to the medium (HCM, Lonza) of Matrigel-retrieved organoids and incubated for given time points at 37°C. For LNP transduction of rescuing mRNAs, organoids were incubated for 90 min with the respective LNP in HCM media with 10 µM RHO-Kinase Inhibitor (Tocris) on a thermal shaker (37°C, 450rpm) prior to the last differentiation step and further cultivated in undiluted GelTrex in HCM medium for additional 72h with or without GW4064 (Sigma-Aldrich) stimulation prior to the assays.

### RNA preparation, reverse transcription, pre-amplification, and PCR analysis

Total RNA was extracted and purified using the AmoyDx FFPE DNA/RNA Kit, (Amoy Diagnostics Co.) according to the manufacturer’s instructions. 100ng of RNA was reverse transcribed using the High Capacity cDNA Reverse Transcription Kit (Applied Biosystems). After pre-amplification, qPCR was carried out with different TaqMan™ Gene Expression Assays (Applied Biosystems). Relative quantification of mRNA was performed according to the comparative 2^-ΔΔCT^ method with SDHA as an endogenous control (see SI for detailed information).

### Structure modeling and molecular dynamics simulations

To analyze the impact of the variant T296I, the ligand binding domain (LBD) structure of FXR (Q96RI1-1, residues 248 to 476) was modeled based on the chenodeoxycholic acid (CDCA)- and NCoA2 peptide-bound X-ray crystal structure of the FXR LBD (PDB ID 6HL1)^16^, representing the active state of FXR, using SWISS-MODEL^24^. To model the inactive state with H12 not interacting with the LBD core, the loop between helix 11 and H12 was remodeled within PyMOL (Schrödinger, LLC, New York). In detail, residues 460 to 466 (^460^VNDHKFT^466^) were removed and readded, pointing away from the LBD core, followed by the α-helix H12.

MD simulations were performed for both the active and inactive states in the presence of the endogenous ligand CDCA and a short peptide sequence from the NCoA2 protein (sequence KENALLRYLLDKD), containing the signature motif LXXLL for binding to an NR^17^. The structural models were prepared for molecular dynamics (MD) simulations using the AMBER21 package^25^. Overall, four different systems were prepared: FXR wildtype in the active state (hereafter termed “active WT”), FXR T296I variant in the active state (“active T296I”), FXR wildtype in the inactive state (“inactive WT”), and FXR T296I variant in the inactive state (“inactive T296I”). Postprocessing and analysis of the MD trajectories were performed with CPPTRAJ^26^ implemented in AmberTools21^25^. For further details, please see the SI Methods.

### Statistical Analysis

Significance tests were performed using the Mann-Whitney U test or Student’s t-test as indicated in the figure captions. The indicated significance levels are n.s. (not significant), *: p ≤ 0.05, **: p ≤ 0.01, ***: p ≤ 0.001, ****: p ≤ 0.0001.

## Results

### The T296I variant is located within the LBD of FXR

The ligand binding domain (LBD) of FXR is critical in regulating the protein’s activity. Residue 296 is located on helix 3 with its side chain facing toward the AF2 interaction surface formed partly by H12 (Fig. 1A and B). We thus hypothesized that variant T296I impacts FXR’s ability to transition from the inactive to the active state. Accordingly, we investigated the effect of variant T296I in MD simulations starting from one of the four configurations: FXR WT in the active state (“active WT”), FXR T296I variant in the active state (“active T296I”), FXR WT in the inactive state (“inactive WT”), and FXR T296I variant in the inactive state (“inactive T296I”) (Fig. 1C). All systems contained the LBD of FXR, the agonist chenodeoxycholic acid (CDCA)^27,28^, and a short peptide sequence of the nuclear receptor coactivator 2 (NCoA2). This setup was chosen to drive the inactive state towards the active state, since the presence of agonists and coactivators has been shown to stabilize the active state^16^. The inactive state was created from the active state through repositioning of the loop region between helix 11 and H12 such that H12 pointed away from the LBD core and had a distance > 45Å to it (distance in the active state 16Å). The setup allowed us to study if the substitution impacts the active state and/or the transition from the inactive to the active state. The ICP-associated M173T variant is located in the DNA-binding domain of FXR and, accordingly, cannot be analyzed with this MD simulations setup.

**Fig. 1:**
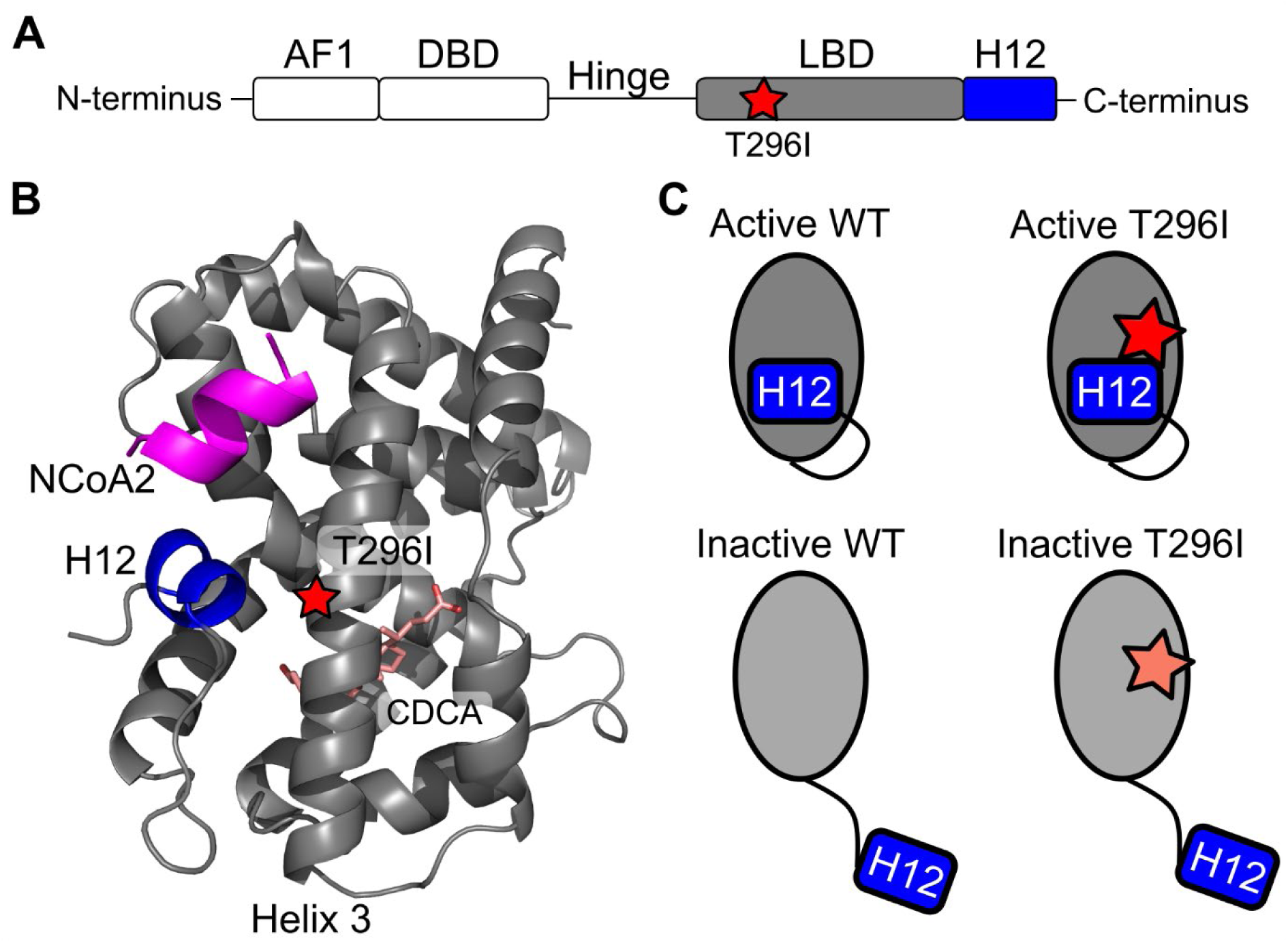
Overview of the protein structure of FXR and the variant site within the LBD. A) Schematic of the domain arrangement of the FXR protein. The N-terminal activation function 1 (AF1) motif is followed by the DNA-binding domain (DBD), which is connected via a flexible hinge region to the LBD and the C-terminal H12. The variant T296I (red star) is located within the LBD. B) Protein structure of the LBD of FXR. The protein systems were modeled based on the crystal structure of agonist-bound FXR LBD^16^ and used for MD simulations, containing additionally CDCA as ligand (pink, shown as sticks) and a short peptide of NCoA2 (magenta). H12 is highlighted in blue and helix 3, containing the variant site T296I (red star), is labeled. The close distance of position 296 to H12 in the active position suggests a potential impact on the conformational change from the inactive to active state. C) Overview of the four systems used as input to MD simulations to study the variant’s impact on the active and the inactive state of the LBD. To differentiate between the different systems, we have consistently used the following color scheme: active WT in darker grey, inactive WT in lighter grey (corresponding to the depicted color of the LBD), active T296I in red and inactive T296I in faded red (corresponding to the depicted color of the star indicating the variant position).

### T296I decreases the transcriptional activity of FXR

To determine the consequences of the FXR missense variant T296I on expression, subcellular localization and target gene induction, human FXRα1 and FXRα2 were cloned from human liver and co-transfected with RXRα into HEK293 cells. Both wildtype (WT) as well as the missense variant were detected within the nucleus of transfected cells (Fig. 2A). Furthermore, protein amounts as determined by western blotting were similar in WT and T296I transfected cells (Fig. 2B). For functional analysis, we used a luciferase expression vector containing the BSEP promoter sequence (BSEP^prom^-Luc.), which was co-transfected with combinations of RXRα and either FXRα1^WT^, FXRα2^WT^, FXRα1^T296I^, or FXRα2^T296I^. The variant effect was further analyzed in transfections without the heterodimer partner RXRα (Fig. S1). The cells were stimulated with an RXR ligand (9-cis-RA, 1μM) and the endogenous FXR agonist CDCA at different concentrations (Fig. 2C, D; Fig. S2A, B). We further validated if the absence of the sodium taurocholate cotransporting polypeptide (NTCP) impacts assay results by additionally co-transfecting cells with NTCP (Fig. S2A). Our results indicate that the presence of NTCP does not alter the overall variant impact, in line with evidence that passive diffusion is the main factor for CDCA uptake into hepatocytes^29^.

**Fig. 2:**
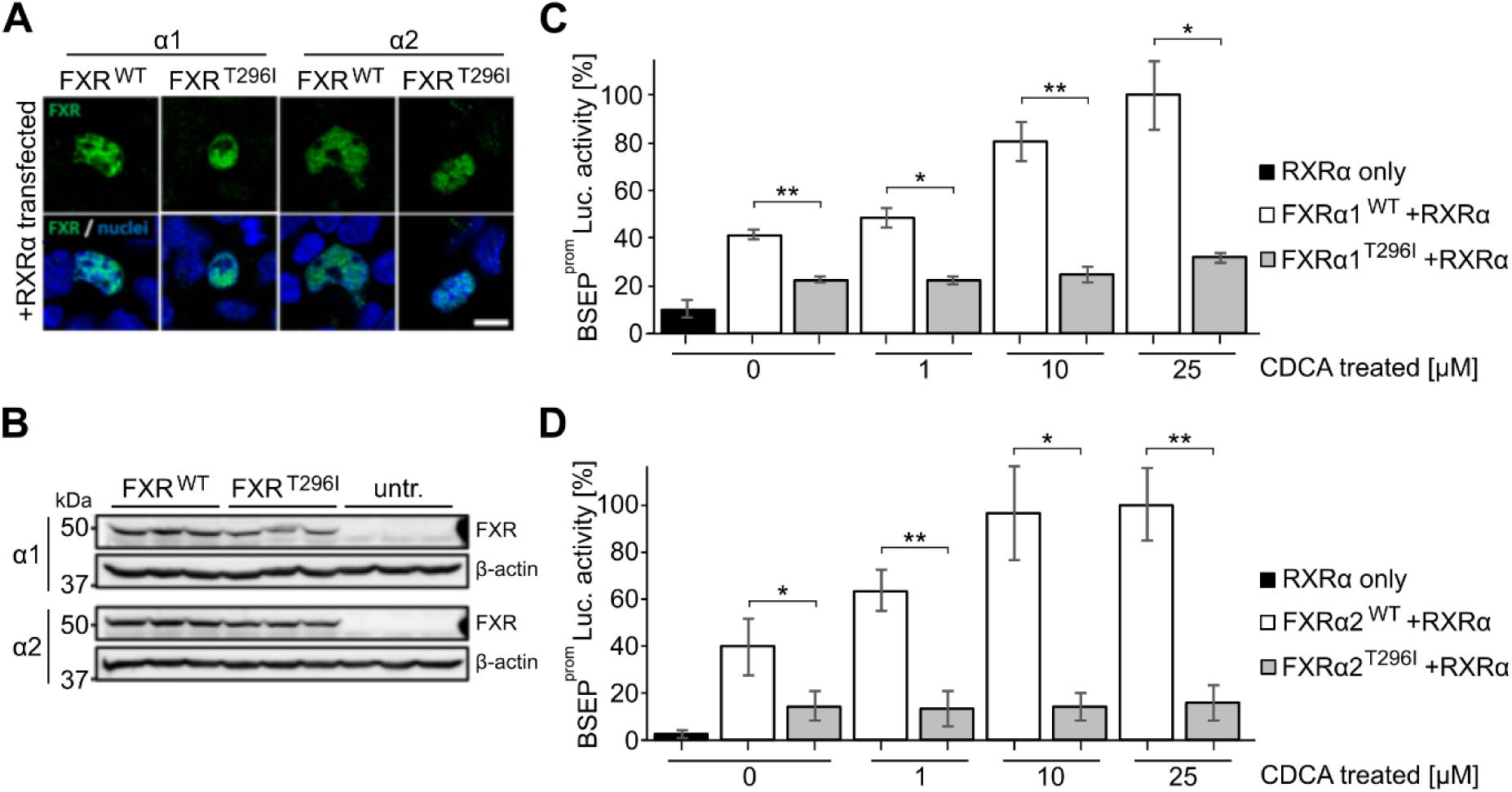
FXR T296I reduces transcriptional activity of the BSEP promoter in transfected HEK293 cells. A) HEK293 transiently co-transfected with RXRα and either FXRα1^WT^, FXRα2^WT^, FXRα1^T296I^, or FXRα2^T296I^ showed correct nuclear localization of the wildtype (WT) and mutant protein. Bar: 10µm. B) Western blot analysis revealed similar protein amounts in cells transfected with the different FXR cDNA constructs, untr.: untransfected controls. C) Analysis of luciferase enzymatic activity after transfection of HEK293 cells with a luciferase reporter gene downstream of the BSEP promoter (BSEP^prom^-Luc) as well as different combinations of RXRα and either FXRα1^WT^ or FXRα1^T296I^, as indicated on the x-axis. Cells were stimulated with the FXR ligand CDCA at indicated concentrations and the RXR ligand 9-cis-RA (1µM). Values were obtained from three independent experiments, in which each condition was tested in triplicates. Values on the y-axis represent the mean and SD, expressed as % luciferase activity. The asterisks indicate a significant difference analyzed by a two-tailed Student’s t-test. D) Analysis of luciferase enzymatic activity after transfection of HEK293 cells with RXRα and either FXRα2^WT^ or FXRα2^T296I^, as indicated on the x-axis. Further conditions were the same as for the FXRα1 isoform in panel C.

Luciferase activity showed an increase in response to transfection with FXRα1/2 and additionally a dose-dependent transactivation of BSEP in response to CDCA^30^. The highest BSEP transactivation was observed when both RXR and WT FXRα1/2 were co-transfected, respectively and cells were stimulated additionally with 9-cis-RA and CDCA 25μM, which was accordingly represented as 100% luciferase activity (Fig. 2C,D). This is in line with previous data showing that the BSEP promoter is regulated by the FXR/RXR heterodimer^31^. The T296I variant resulted in a reduced transcriptional activity in both FXR isoforms (Fig. 2C, D) independent of the used CDCA concentration (0, 1, 10, or 25µM). T296I showed a significant decrease of basal BSEP transactivation with 22.5% and 14.7% for the FXRα1 and FXRα2 isoform, respectively, compared to 41.1% and 39.7% for the FXR WTα1/α2 isoforms in the absence of CDCA. Basal activity for both FXR isoforms has been identified previously^30,32^. At 1µM CDCA treatment, the T296I variant reached 22.1% and 13.1% of luciferase activity while the wildtype FXRα1 and FXRα2 isoforms showed significantly higher luciferase activity at 48.4% and 63.6%, respectively. At a dose of 10µM CDCA the variant reached 24.9% and 14.3% of luciferase activity, while the FXR WT isoforms showed responses with 80.4% and 96.4%, respectively. At 25µM CDCA, the variant showed significantly reduced luciferase activity of 31.6% and 16.0%, compared to the maximal response of FXR WTα1/α2, set to 100% for each isoform. The activity of the T296I variant increased with higher ligand concentration (Fig. S2C), supporting the hypothesis that the variant did not impair ligand binding (Fig. S7, S8 and S9). Furthermore, the T296I variant reduced transcriptional activity of the SHP promoter, another FXR target (Fig. S1B, S2D).

In summary, even though the subcellular localization and protein expression levels were unaffected, the presence of the T296I missense variant resulted in a significant decrease in the transactivation of FXR target genes *in vitro*.

### FXR T296I leads to an increased distance between H12 and the substitution site, indicative of a less favorable active state

Using MD, we investigated the molecular mechanisms underlying the decreased activity of the T296I variant. All simulation systems showed minor structural variability with respect to the binding of CDCA and NCoA2 and the FXR LBD structure up to and including helix 11 (Fig. S3). Based on the crystal structure of the FXR LBD^16^, representing the agonist-bound active state, the WT T296 likely interacts with T466 preceding H12 (Fig. 3A). Accordingly, we measured the distance between residue T296 and T466 during MD simulations and compared it to the reference distance in the crystal structure. For the active states, the variant showed an increase in the distance (Fig. 3B). We determined the frequency of occurrence when the reference cutoff distance was reached (Fig. 3C, Table S1). Across the 15 replicas, the active WT system was found in approx. 27% of the time in the active state according to distances below the reference distance. The frequency of occurrence was significantly lower for active T296I (0.40%). The inactive systems showed initially high distance values, as expected. Inactive WT reached below the reference distance in 6 out of 15 replicas and often stayed within this active state for the remainder of the simulation time, indicating that the active state is the preferred one under the simulation conditions (Fig. S4, Table S1). Inactive T296I only reached the reference distance in one replica (Fig. S4, Table S1) and, accordingly, the frequency of reaching the reference value was significantly reduced in the inactive T296I system compared to the inactive WT system (Fig. 3C).

**Fig. 3:**
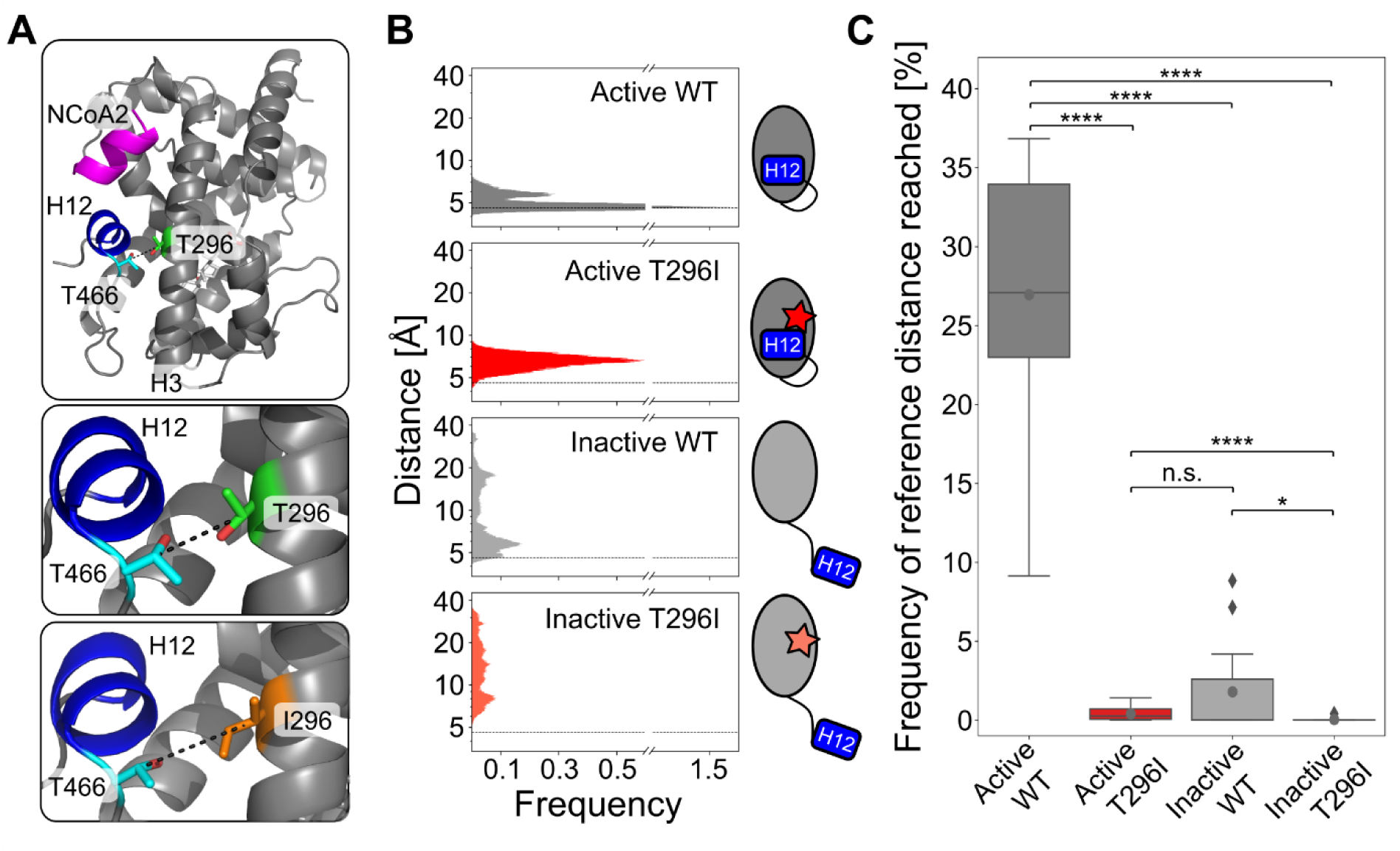
Distance analysis between residues T296 and T466 over MD simulation time. A) Depiction of the distance measured within the LBD of FXR between the C_β_ atoms of residue T296 and T466. The mean distance is increased in the active T296I (6.6Å) compared to the active WT (5.0Å). B) Histograms of measured distances over all 15 MD runs (see Fig. S4) for each analyzed state. The reference distance (4.6Å) as measured in the agonist-bound crystal structure^16^ is depicted as dashed lines. C) Frequency of occurrence that the respective system is in the active state. For each replica (Fig. S4), the percentage of reaching a distance below the reference distance (4.6Å) is depicted as a boxplot. Individual values are shown in Table S1. Boxes depict the quartiles of the data with the median (straight black lines) and mean (grey dots) indicated; the whiskers indicate the minimum and the maximal values, outlier points are depicted as rhombus. Differences in the mean values were statistically evaluated using a two-sided Mann-Whitney U test (*N* = 15, n.s.: not significant; *: *p* ≤ 0.05, ****: *p* ≤ 0.0001).

Overall, we observed an increase in the distance between residue T296 and T466 for the T296I variant in the active state. In line, for systems started either from the active or inactive state, T296I led to a significant decrease in the frequency of occurrence in reaching the reference distance compared to the WT. This data indicates that the active state is destabilized in the variant.

### The correct positioning of H12 in the active state is reduced in the T296I variant

As the correct positioning of H12 within NRs is crucial for activity^33–36^, we visually analyzed the simulation trajectories of the inactive systems and exemplarily show the conformational change from the inactive to the active state for one out of several MD replicas of the inactive WT system showing this transitioning (Fig. 4A and Movie S1). For the inactive T296I, we show the MD replica where H12 positioning was closest to the conformation in the active state (Fig. 4B and Movie S2). Further, we analyzed the impact of T296I on the positioning of H12 using the root mean square deviation (RMSD) of residues 466 to 473 (H12 and preceding T466) over the MD simulation time (Fig. 4C and D). As a reference state, we employed the active conformation based on the crystal structure of the agonist-bound FXR LBD^16^. For the active WT, a skewed Gaussian function was fitted to the RMSD histogram, yielding a mean of 1.9Å (Fig. S5 and Fig. 4C). Over the entire MD simulations, more than half of the time (∼56%, Table S2 and Fig. 4D) the active WT had an RMSD below this mean value, which we used as a further reference to indicate reaching the active state. Comparing the active systems revealed a significant shift in the distributions, with a larger mean of active T296I indicating a higher deviation from the active positioning (active WT mean: 1.9Å, active T296I mean: 2.6Å, *p* = 0.0022, two-sided *t*-test, Fig. S5). The frequency of reaching the active state was significantly lower for active T296I, inactive WT, and inactive T296I systems compared to active WT (Fig. 4D). While the inactive WT reached the reference RMSD value in four out of 15 replicas, the inactive T296I reached it in one replica (Fig. S6). Furthermore, while the inactive T296I did not stay in the active state long (frequency: 0.01% in replica no. 6), the inactive WT – once reaching the active state – showed often prolonged persistence times (frequency: 15.40% in replica no. 2, 0.26% in replica no. 6, 10.17% in replica no. 8, 14.29% in replica no. 13) (Fig. S6 and Table S2). The comparison between inactive WT and inactive T296I indicated a similar trend as observed for the distance analysis but did not reach the significance level (Fig. 4D).

**Fig. 4:**
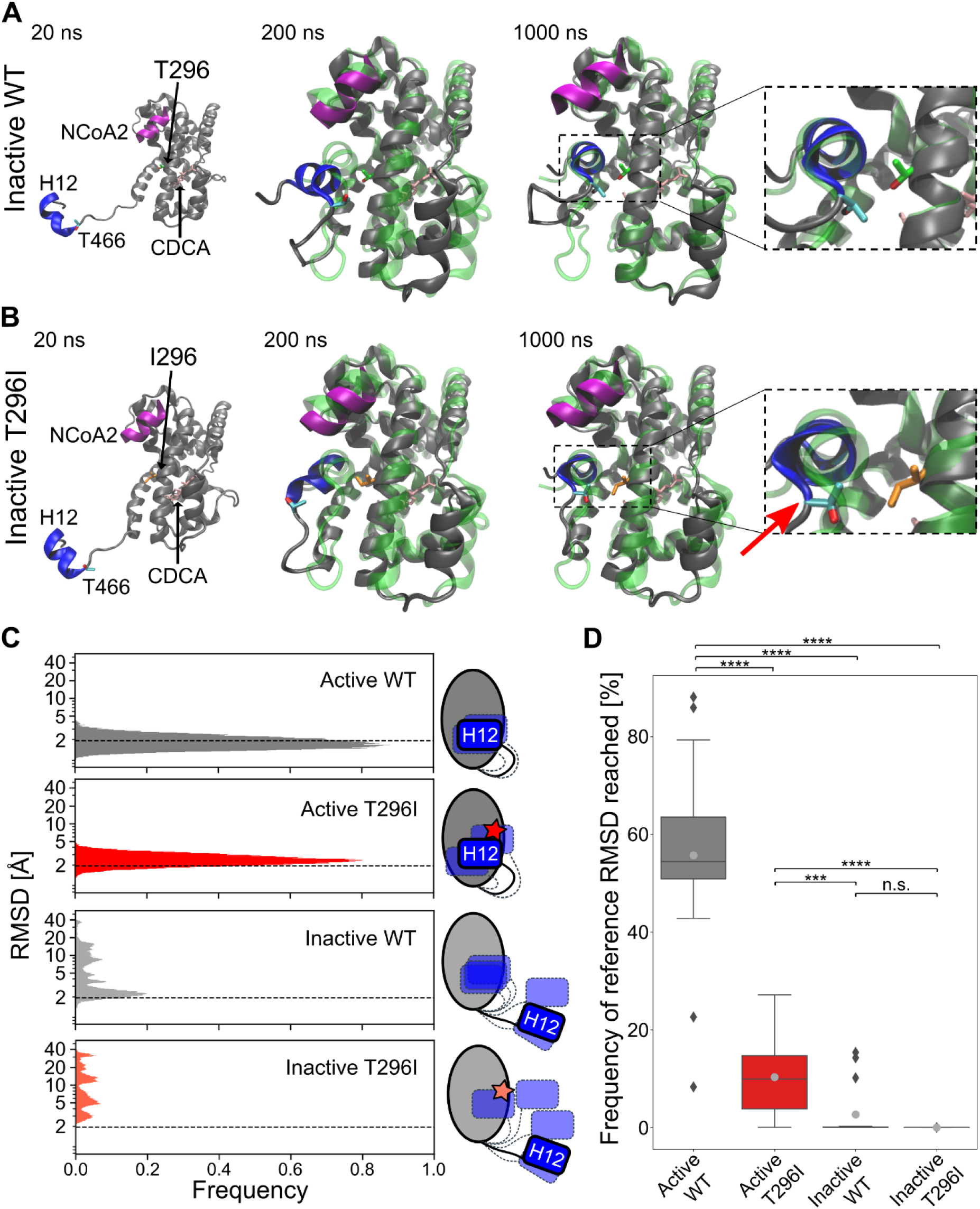
Positioning of H12 during MD simulations. A) Conformational transitioning of the inactive WT (replica no. 2, Movie S1, Fig. S4 and S6) over the MD simulation time. The initial active state (based on the crystal structure of agonist-bound FXR LBD^16^ is depicted as a green, translucent cartoon structure. B) Transitioning of the inactive T296I (replica no. 6, Movie S2, Fig. S3 and S5) over the MD simulation time. The initial active state is depicted as a green, translucent cartoon structure. Important residues (variant site 296 [green or orange, shown as sticks], T466 [cyan, shown as sticks], H12 [blue], NCoA2 [magenta], and bound ligand CDCA [pink, shown as sticks]) are additionally depicted in A and B. After 1000ns, H12 of the inactive WT showed good alignment with the active structure, while H12 of the inactive T296I showed structural deviations (red arrow). C) Histograms of RMSD values of H12 and the preceding T466 over all 15 MD replicas for each analyzed state. Data for each MD replica are depicted in Fig. S5. A skewed Gaussian function was fitted to the distribution of active WT (Fig. S5), and the obtained mean (1.9Å) was used as a reference cutoff (dashed lines). D) Frequency of occurrence a system spends in the active state, i.e., when the reference cutoff is reached. For each replica (Fig. S6), the frequency of occurrence was determined and depicted within boxplots. Individual values are shown in Table S2. Boxes depict the quartiles of the data with the median (straight black lines) and mean (grey dots) indicated; the whiskers depict the minimum and the maximum values, outlier points are depicted as rhombus. Differences in the mean values were statistically evaluated using a two-sided Mann-Whitney U test (*N* = 15, n.s.: not significant; ***: *p* ≤ 0.001, ****: *p* ≤ 0.0001).

Overall, H12 positioning is significantly structurally deviating with respect to the reference active state for all systems compared to the active WT. While the active T296I could reach the reference cutoff, it did so for a significantly decreased amount of time compared to the active WT, again indicating that the active conformation is less favorable in the variant. Although inactive WT and inactive T296I could both reach the cutoff, the inactive WT reached it more frequently and for a longer time. However, the differences to inactive T296I are not significant. Our data indicate that unbiased MD simulations on the μs-scale can sample the transition from the inactive to the active state (see Table 1) and that this conformational change is less frequent in the variant.

**Table 1:** Overview of the MD simulation results.

|  | Distance criteria | RMSD criteria |
| --- | --- | --- |
| Inactive WT reaching active state | 6 out of 15 runs | 4 out of 15 runs |
| Inactive T296I reaching active state | 1 out of 15 runs | 1 out of 15 runs |
| Significance between WT and T296I<sup>a</sup> | * ($p = 0.016$) | n.s. ($p = 0.076$) |

<sup>a</sup> Using one-sided *t*-test.

### The T296I variant is associated with reduced expression of FXR target genes

To investigate the mRNA expression of FXR and two of its targets (*ABCB11* encoding for BSEP and *NR0B2* encoding for SHP; we have used the protein names in the following for coherence throughout the manuscript) in the PFIC patient carrying the T296I variant, we performed qPCR analysis using FFPE samples from the patient’s liver taken at the time of transplantation. Additionally, FFPE-liver samples from two cirrhotic adult patients and a healthy adult control were included in the analysis. To address the putatively low RNA integrity after isolation from FFPE samples, we employed TaqMan Gene Expression Assays targeting different regions of the FXR and BSEP transcripts. While FXR mRNA expression was similar between the patient and control and the two cirrhosis livers (Fig. 5A), BSEP mRNA and SHP mRNA expression was strongly reduced in comparison to the healthy control but also the two cirrhotic liver samples (reduction to 3.03, 1.82 and 10.91% of healthy control for BSEP Taq1, BSEP Taq2, and SHP, respectively) (Fig. 5B). These findings clearly suggest that our *in vitro* and *in silico* data authentically reflect the impaired transcriptional target gene activation by the FXR T296I variant observed in the patient’s liver specimen.

**Fig. 5:**
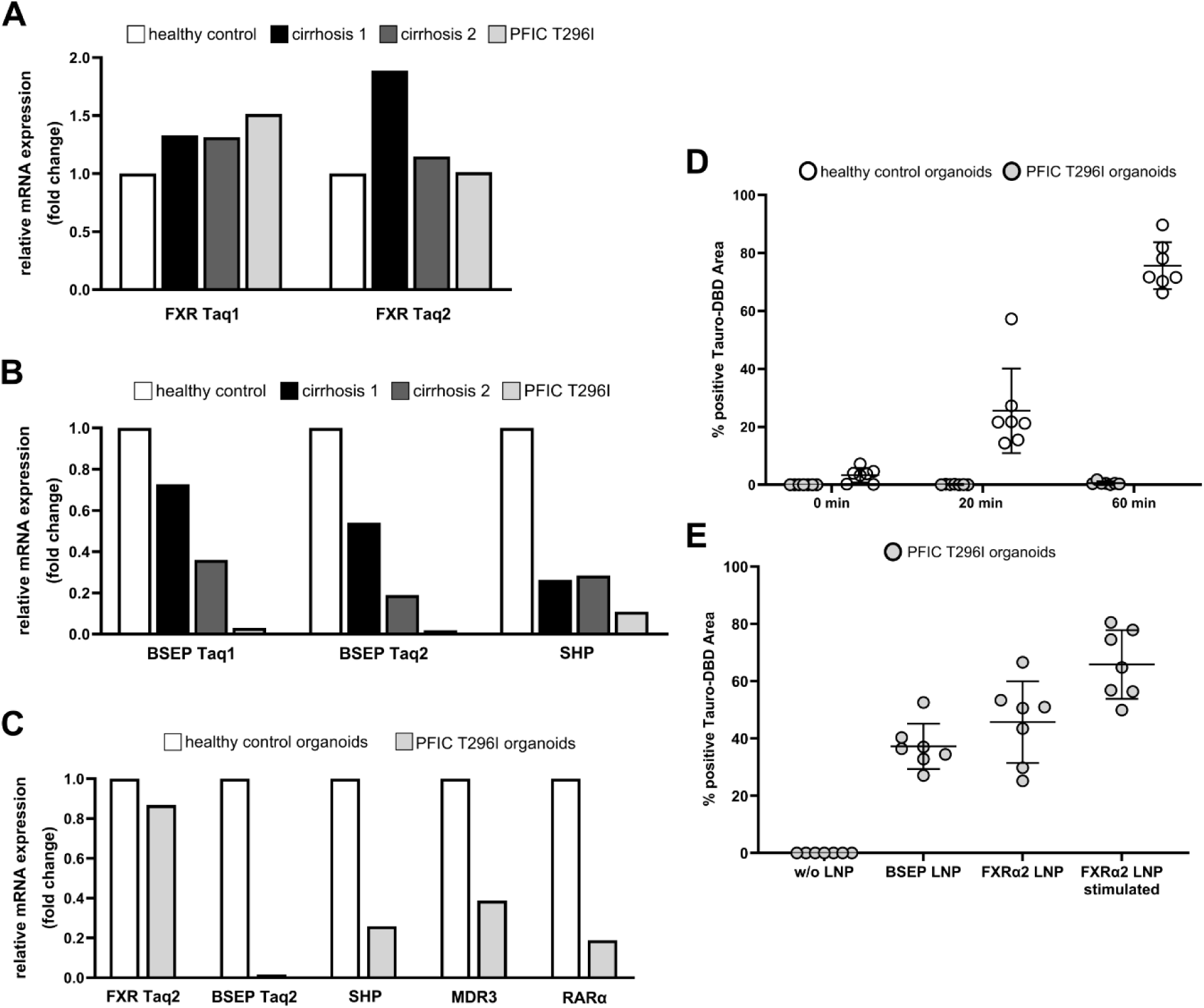
FXR, BSEP, and SHP expression in the patient’s liver tissue and patient’s iPSC-derived hepatic organoids. A) Relative mRNA expression of FXR using two different TaqMan probes (Taq1 and Taq2) showed similar levels in the liver of a healthy control (white bars), two samples from cirrhotic livers (black bar and dark grey bar) as well as the patient (light grey bar). B) FXR target gene expression of BSEP and SHP were lower in the two cirrhotic livers as compared to the healthy control but were further reduced in the sample from the T296I PFIC patient. C) Patient’s iPSC-derived organoids exhibit FXR expression and marked reduction of target gene expression (BSEP, SHP, MDR3, RARα). D) Functional assessment of BSEP-mediated transport of Tauro-nor-THCA-24-DBD (Tauro-DBD) exhibited proper time-dependent transport (0min - 60min) in control iPSC-derived hepatic organoids but absence in the patient’s iPSC-derived organoids. E) Delivery of BSEP or FXRα2 (± stimulation with GW4064) mRNA via lipid nanoparticles (LNP) restored Tauro-DBD transport.

To get more functional insights, we reprogrammed peripheral blood mononuclear cells from the FXR T296I patient into induced pluripotent stem cells (iPSCs), which were subsequently differentiated into hepatic organoids. qRT-PCR of these patient-iPSC-derived liver organoids revealed an uncompromised expression of FXR (86.8% of healthy control), but a marked reduction of mRNA expression of FXR target genes BSEP, SHP, MDR3, and RARα (1.6, 25.8, 38.7, and 18.8%, respectively) (Fig. 5C). As the hepatic organoids build apical luminal structures resembling bile canaliculi, we were able to study the transport of the fluorescent BSEP-specific substrate Tauro-nor-THCA-24-DBD (Tauro-DBD) into such luminal cavities. After recovery of hepatic organoids from the Matrigel domes, the organoids were incubated with the fluorescent substrate for 0min, 20min, or 60min at 37°C and the proportion of fluorescence-positive luminal structures over the total area of luminal structures was assessed by fluorescence microscopy and subsequent image analysis. Control organoids exhibit intact Tauro-DBD transport capabilities (3.3% at 0min, 25.6% at 20min, and 75.6% at 60min Tauro-DBD-positive area; average over seven independent measurements), while organoids obtained from the T296I-mutant iPSCs did not show Tauro-DBD transport (<0.1%, <0.1% and 0.6% at the respective timepoints) (Fig. 5D). Lipid nanoparticle (LNP)-mediated mRNA delivery allows gene supplementation during the formation of hepatic organoids. We were able to restore BSEP-mediated Tauro-DBD transport after LNP-based delivery of BSEP-mRNA or FXRα2-mRNA in the patient’s iPSC-derived organoids to levels of 37.2% and 45.7%, respectively (Fig. 5E). Stimulation of FXR activity by 1µM GW4064 resulted in an increased BSEP transport activity (65.8%), suggesting that FXR-mediated signaling is authentically reflected in this iPSC-derived organoid model. This strengthens our data that the FXR T296I variant fails to activate BSEP gene expression, thus, causing a PFIC phenotype.

## Discussion

In this study, we combined computational studies with *in vitro* experiments to analyze the impact of the PFIC-associated *NR1H4* T296I variant on FXR protein expression, subcellular localization, and function. While the introduction of the missense variant into the α1 or α2 FXR isoform did not affect protein expression and nuclear localization *in vitro*, it significantly reduced activation of the FXR target genes BSEP and SHP. A strong reduction of BSEP and SHP was also observed in the patient’s liver at the time of transplantation and in iPSC-derived hepatic organoids of the patient, while FXR expression levels were similar to controls. Partial rescue of the impaired BSEP-mediated Tauro-DBD transport in T296I organoids was achieved using LNP with BSEP or FXRα2 WT mRNA, further validating that the variant is causative of the reduced BSEP expression and subsequent phenotype. Using our computational approach, we elucidated a detailed mechanism for the effect of the variant on the conformational transition of the LBD from the inactive to the active state. The variant showed a significantly reduced tendency to reach the active state, which can explain the *in vitro*-identified decreased target gene expression and thus the PFIC phenotype of the patient^11^. Employing MD simulations for an atomistic analysis of this variant indicated a molecular mechanism underlying the protein dysfunction and showcases implications for general FXR protein activation.

We describe conformational changes from an inactive to the known active state of the FXR LBD in unbiased MD simulations. To drive the system towards the active state, we used a coactivator peptide and the most potent *in vivo* agonist CDCA^37^ within the MD systems as both coactivator peptide and ligand binding have been shown to induce and stabilize the active state *in vitro*^16^. Depending on the analysis, the inactive WT system reached the active state in 27% or 40% (4/15 replicas for H12 RMSD analysis and 6/15 replicas for distance analysis, respectively) of the simulations (Table 1). A dynamic movement from the inactive to the active state (and potentially reverse, at least in ligand-free states) may occur in the nanosecond time scale as indicated by time-resolved fluorescence anisotropy decay studies on PPARγ^35^. Chrisman et al. showed conformational changes of the H12 within the NR PPARγ towards an almost-active state within the µs to ms time scale range in unbiased MD simulations^15^. Within our MD simulations, the LBD of FXR might sample conformational spaces usually not available due to sterical hindrances either by the not considered parts of FXR (DNA binding domain and linker sequences) or due to hetero-dimer binding partners. Thus, it is not surprising that in several replicas, H12 did not move into the active position within the 1µs of simulation time due to being trapped in other energy minima. Still, fast transitioning from the inactive to the active state was observable in some replicas. Comparing the difference between inactive WT and inactive T296I systems in reaching the active state, we observed a decrease by a factor of 3.8 to 5.7 (inactive WT: 26.7% or 40%; inactive T296I: 7%). This is in good agreement with the transcriptional activity reduction of T296I compared to WT as shown in the luciferase assay for BSEP and SHP as well as the patient’s liver tissue (Fig. 2, Fig. 5). Note that we have not investigated the impact on co-regulator recruitment or other FXR functions. These limitations may be addressed through more detailed investigations, which are beyond the scope of this study but present valuable opportunities for future research.

Overall, the variant T296I, while also impacting the active state, likely exerts its negative impact on protein activity due to a change in the structural dynamics of the inactive-to-active state transition. Our results indicate that the T296I protein does not reach the active state fully and less frequently compared to the WT protein. Furthermore, from the analysis of the MD simulations, insights into the activation mechanism of the LBD were gained. The presence of the ligand and co-activation peptide allows FXR to switch into an active state and stably stay within this state. This is in line with previous NMR or MD studies in RORγ^14^, PPARγ^15,38^, and FXR or FXR/RXR heterodimers^19–21^. By contrast, the transition from inactive to active FXR has previously not been observed in MD simulations. Our setup of the MD simulations can be useful to predict the impact of other missense variants on FXR function and potentially strengthen studies on FXR targeting, enabling detailed evaluations of the molecular mechanism of drugs based on their impact on the activation transitioning.

## Abbreviations

FXR: farnesoid X receptor
LBD: ligand binding domain
PFIC: progressive familial intrahepatic cholestasis
BA: bile acids
RXR: retinoic X receptor
MD: Molecular Dynamics
NR: nuclear receptor
RMSD: root mean square deviation
H12: helix 12
AF: activation function
OCA: obeticholic acid
CDCA: chenodeoxycholic acid
NCoA2: nuclear receptor coactivator 2

## Funding

This study was supported by the BMBF through HiChol (01GM1904A and 01GM2204A to V.K. and C.D.; 01GM1904A and 01GM2204B to H.G.; 01GM1904B and 01GM2204C to U.B., A.S., E.P., T.C.).

## Acknowledgment

Expert technical assistance by Paulina Philippski and Aileen Nötzold is gratefully acknowledged. We are grateful for computational support by the “Zentrum für Informations und Medientechnologie” at the Heinrich-Heine-Universität Düsseldorf. We thank the Gauss Centre for Supercomputing e.V. (www.gauss-centre.eu) for funding the project [project IDs: VSK33; FIC1; FXR] by providing computing time through the John von Neumann Institute for Computing (NIC) on the GCS Supercomputer JUWELS^1^ at Jülich Supercomputing Centre (JSC).

## Conflict of interest statement

The authors declare that there is no potential conflict of interest.

## Notes

### Competing Interest Statement

The authors have declared no competing interest.

### Summary of Updates

- We have extended our luciferase-based cellular assays to analyze different OCA concentrations as well as the endogenous bile acid CDCA. We included this data, in addition to data for the ICP-associated variant M173T (van Mil et al. 2007), in the main text and the SI. - We have added a detailed investigation of the variant's impact in iPSC-derived hepatic organoids, derived from the patient carrying the T296I as well as a control. We have further added MDR3 and RARα to the panel of investigated genes and all FXR-target genes showed a marked downregulation while the FXR expression is comparable to control levels, further strengthening the data. Additionally, investigating the functionality of BSEP using the fluorescent BSEP-specific substrate Tauro-nor-THCA-24-DBD (Tauro-DBD) showed a drastic reduction of BSEP-mediated transport in the case of the T296I variant. Of note, this phenotype could be rescued with lipid nanoparticle-mediated mRNA delivery of BSEP or WT FXRα2, verifying that the variant is the main driver for the observed phenotype. This data has been integrated into the updated Fig. 5, and the description of the materials and methods used for the organoid generation and investigation has been added to the main text and SI.

