## Supporting Information for "Impaired transitioning of the FXR ligand binding domain to an active state underlies an FXR-associated PFIC phenotype"

### Supplementary Results

#### FXR T296I reduces transcriptional activity of the BSEP and SHP promoter

We further analyzed the impact of the FXR T296I variant alone as well as in combination with RXRα in HEK293 cells. First, we used a luciferase expression vector containing the BSEP promoter ($BSEP^{prom}$-Luciferase), which was transfected with either $FXR\alpha1^{WT}$, $FXR\alpha2^{WT}$, $FXR\alpha1^{T296I}$, or $FXR\alpha2^{T296I}$ alone and in combination with RXRα (Fig. S1A). The highest BSEP transactivation was observed when RXRα and $FXR\alpha1^{WT}$ or $FXR\alpha2^{WT}$, respectively, were co-transfected, which was accordingly represented as 100% luciferase activity. Transfection of $FXR\alpha1^{WT}$ or $FXR\alpha2^{WT}$ alone resulted in luciferase activity of 19.2% and 34.4%, respectively. However, when $FXR\alpha1^{T296I}$ or $FXR\alpha2^{T296I}$ was transfected alone, there was a significant decrease in luciferase activity to 11.7% and 4.9%, respectively. Similarly, the co-transfection of RXRα with the $FXR\alpha1^{T296I}$ or $FXR\alpha2^{T296I}$ led to a significant decrease to 26.1% and 13.1%, respectively, when compared to the RXRα/WT FXR co-transfection. In conclusion, the T296I variant resulted in a significant decrease of BSEP transactivation both in single transfections with FXR as well as in co-transfections with RXRα, indicating that the variant impacts protein activity independently of its function as a heterodimer with RXRα or as a monomer.

For further functional analysis, we used a luciferase expression vector containing the SHP promoter sequence ($SHP^{prom}$-Luciferase), which was co-transfected with several combinations of RXRα and either $FXR\alpha1^{WT}$, $FXR\alpha2^{WT}$, $FXR\alpha1^{T296I}$, or $FXR\alpha2^{T296I}$. The highest SHP transactivation was observed when both RXR and $FXR\alpha1^{WT}$ or $FXR\alpha2^{WT}$, respectively, were co-transfected, which we represented as 100% luciferase activity (Fig. S1B). Transfection of $FXR\alpha1^{WT}$ or $FXR\alpha2^{WT}$ alone resulted in luciferase activity of 17.7% and 18.5%, respectively, in comparison to co-transfection with both RXRα and $FXR\alpha1^{WT}$ or $FXR\alpha2^{WT}$. However, when the $FXR\alpha1^{T296I}$ or $FXR\alpha2^{T296I}$ variant was transfected alone, there was a significant decrease in luciferase activity to 7.4% and 9.5%, respectively, when compared to $FXR\alpha1^{WT}$ or $FXR\alpha2^{WT}$. Similarly, the co-transfection of RXRα with $FXR\alpha1^{T296I}$ or $FXR\alpha2^{T296I}$ led to a significant reduction in luciferase activity, with decreases to 43.7% and 71.7%, respectively, in comparison to the co-transfection of RXRα and $FXR\alpha1^{WT}$ or $FXR\alpha2^{WT}$. In conclusion, the presence of T296I resulted in a substantial decrease in SHP transactivation in the transfected cells.

**FXR T296I reduces transcriptional activity more strongly than FXR M173T and independently of the ligand type and concentration used for stimulation**

To stimulate FXR we used both the endogenous ligand CDCA as well as the semisynthetic CDCA-derivative obeticholic acid (OCA)[1] (10µM) (see Fig. S2). We further investigated the impact of the T296I variant in comparison to the milder and ICP-associated variant M173T[2], which enables an assessment of the range of variant impacts. M173T showed reduced transcription of the downstream FXR targets BSEP and IBABP in luciferase assays while DNA binding and heterodimerization to RXR was unimpaired[2]. Due to its localization in the DNA-binding domain of the FXR protein, we could not investigate the M173T variant effect using the MD simulations setup, as it is focused on the ligand binding domain. We used a luciferase expression vector containing the BSEP promoter (BSEP$^{prom}$-Luciferase) and co-transfected HEK293 cells with RXRα and either FXRα1$^{WT}$, FXRα2$^{WT}$, FXRα1$^{T296I}$, FXRα2$^{T296I}$, FXRα1$^{M173T}$, or FXRα2$^{M173T}$ (Fig. S2). We additionally transfected the cells with the sodium taurocholate cotransporting polypeptide (NTCP)[3] (Fig. S2A). Cells were stimulated with the RXR agonist 9-cis-retinoic acid (9-cis-RA, 1µM) and different concentrations of CDCA (0, 1, 10, or 25µM). The highest transactivation activity was observed using the highest concentration of CDCA in cells transfected with RXRα and FXRα1$^{WT}$ or FXRα2$^{WT}$ and was accordingly set to 100%, respectively. The variant T296I showed a significant decrease in luciferase activity for all CDCA concentrations in both FXR isoforms, compared, respectively, to RXRα and FXRα1$^{WT}$ or FXRα2$^{WT}$ at the same CDCA concentrations. The ICP-associated M173T variant showed higher activity than the T296I variant through all conditions. When compared to FXRα1$^{WT}$, FXRα1$^{M173T}$ showed significantly reduced activity. However, no significant change was found for FXRα2$^{M173T}$ compared to FXRα2$^{WT}$ (Fig. S2A).

Next, we omitted the NTCP transfection to verify that NTCP expression is not necessary for CDCA uptake[4] (Fig. S2B). Again, the T296I FXR variant showed in both isoforms a significant decrease for all used CDCA concentrations (0, 1, 10, and 25µM) when compared to the respective FXR WT conditions. The M173T variant showed a decreased but not significantly different luciferase activity in both isoforms compared to the respective FXR WT conditions for all CDCA concentrations, indicating a milder effect than what the M173T variant showed when NTCP was additionally transfected (Fig. S2A). In conclusion, NTCP expression did not change the effect of CDCA on the T296I variant, while the effect of CDCA on the milder variant M173T may be influenced by the presence or absence of the transporter.

Furthermore, we validated the results using a different FXR agonist, the semisynthetic CDCA-derivative OCA at different concentrations (0, 0.1, 1, and 10µM) (Fig. S2C). The T296I variant showed reduced luciferase activity for FXR isoforms and for all OCA concentrations when compared to the respective FXR WT conditions. The T296I variant showed higher luciferase activity with increasing OCA concentration similar to the FXR WT, indicating that differences in ligand binding do not underlie the reduced activity (see Fig. S7, S8 and S9). The M173T variant showed reduced luciferase activity in the FXRα1 isoform while no significant differences were observed in the FXRα2 isoform when compared to the respective FXR WT conditions.

To further expand the investigation to a different FXR target, we investigated the variant effects on a luciferase expression vector containing the SHP promoter ($SHP^{prom}$-Luciferase) and stimulated cells with different CDCA concentrations (0, 1, 10 and 25µM) (Fig. S2D). The T296I variant showed significantly reduced luciferase activity at higher CDCA concentrations (10µM and 25µM) when compared to the WT FXR conditions, respectively, in the FXRα2 isoform. The M173T variant showed no significant difference compared to the WT FXR conditions, respectively.

**Structural variability during the MD simulations**

The structural variability of the systems was analyzed by root mean square deviations (RMSD) with respect to the first production frame over the MD trajectories (Fig. S3). The results indicate that the CDCA ligand remained in its initial binding mode (Fig. S3A, B) and showed minor structural differences between the active T296I and the inactive WT system (Fig. S3C-E). The LBD of FXR, excluding the flexible region of helix 12 (H12) and the preceding loop, showed a constant RMSD over the simulation time with no significant differences between the four systems (Fig. S3C), further indicating that the LBD without H12 did not undergo significant conformational changes over the simulation time. Expectedly, the inactive WT and inactive T296I systems showed higher structural variability of H12 and the preceding loop region compared to the active systems, indicating higher mobility of this part of FXR in the inactive systems. Furthermore, in the active T296I, the mobility of H12 and the preceding loop region is significantly higher than in the active WT system (Fig. S3D), in line with further analyses of H12 mobility (see Fig. 4). The mobility of the NCoA2 peptide is similarly low in all systems, indicated by generally low RMSD values. Larger values, especially visible in the inactive systems, are indicative of higher mobility, and the displacement of the peptide was also visually observed in several replicas. Except for the comparison of active T296I to inactive T296I, which reveals a significantly increased mobility, the differences between the systems were not significant.

**Geometric analyses of the MD trajectories**

The distance between residue T296, the mutation site, and the threonine preceding H12, T466, was measured over the MD simulation time (Fig. S4). The distance was measured between the $C_\beta$ atoms of both residues to avoid biases due to rotations of the side chains. For each replica, the time the system spent with a distance below the reference distance cutoff (taken from the initial active structure) was calculated (Table S1).

The structural variability of H12 was measured using the RMSD of all atoms of the H12 residues and the preceding T466 ($^{466}$TPLLCEIW$^{473}$). Beforehand, the least mobile part of the FXR LBD was identified, and the trajectories were fitted to this core. As a reference, the initial active WT system was used. The RMSD was determined over the entire MD simulations time; a histogram of the values revealed a skewed Gaussian curve for the active WT and active T296I system (Fig. S5). Since the distribution of active WT can be expected in a physiological, uninhibited, active system, the mean of a fitted skewed Gaussian curve was used as a reference for the active state. The histogram of the active T296I is significantly shifted (assessed by a two-sided *t*-test) towards higher RMSD values compared to the active WT, indicating increased mobility of H12. The RMSD values over the MD trajectories of the individual replicas are depicted in Fig. S6. The inactive WT reached the reference RMSD value of the active WT in four out of 15 replicas, whereas the inactive T296I only reached the reference in one out of 15 replicas. For each replica, the time the system showed an RMSD below the reference RMSD cutoff (mean of the fitted skewed Gaussian curve on active WT) was calculated (Table S2).

**Melting temperature measurement of WT and variant protein in the absence or presence of ligand**

To exclude the possibility that the variant‘s decreased protein activity identified in luciferase assays is due to changed ligand binding, we measured the thermostability of the FXR protein in the presence and absence of the agonist OCA (INT-747). FXR WT and FXR T296I variant proteins were expressed in *E. coli* with a His-tag and SUMO-tag to aid in purification and solubility (Fig. S7). In the absence of ligand, both FXR WT and variant FXR T296I had comparable melting temperatures, indicating that the protein structure is not significantly changed due to the amino acid substitution (Fig. S8 and Fig. S9, Welch’s *t*-test WT+DMSO vs. T296I+DMSO: n.s.). As expected, the presence of the ligand induced a shift in the melting temperature curve and led to a significantly lower melting temperature for the WT FXR protein by 2.32+/-0.70°C (Welch’s t-test WT+DMSO vs. WT+OCA: *). For the variant protein FXR T296I, the presence of the ligand induced a similar shift by 2.78+/-0.49°C (Welch’s t-test T296I+DMSO vs. T296I+OCA: ***) and resulted in a lower melting temperature, comparable to the FXR WT in the presence of ligand (Welch’s t-test WT+OCA vs. T296I+OCA: n.s.). Accordingly, agonist OCA binding to the protein was not disturbed by the variant, in line with previous studies on the NR retinoic-acid related-orphan-receptor-C (RORγ) suggesting that the ligand entry and exit pathway occurs via the so-called “backdoor” pathway and, thus, away from the variant site[5]. Of note, other NRs such as estrogen receptor, androgen receptor, and glucocorticoid receptor can be classified by a different ligand entry via a Helix 3/Helix 7/Helix 11 interface[6], while FXR has been grouped with RORγ and PPARγ[5,7]. Similarly, unbinding MD simulation studies on the FXR LBD indicated egress pathways in line with the backdoor pathway, facing away from the Helix 12 and the variant site, as most favorable for the agonistic ligand GW4064[8].

**Visualization of the conformational change from the inactive to the active state**

The trajectories of the inactive WT replica no. 2 (Movie S1) and the inactive T296I replica no. 6 (Movie S2) were chosen as representative trajectories of the respective system reaching the active state. In the case of inactive T296I, the system moved close to the active state but did not entirely reach it and/or showed higher mobility there. The reference state of the initial active WT is repeatedly shown in the movie as a green translucent representation to aid in the judgment of H12 positioning.

### Supplementary Figures

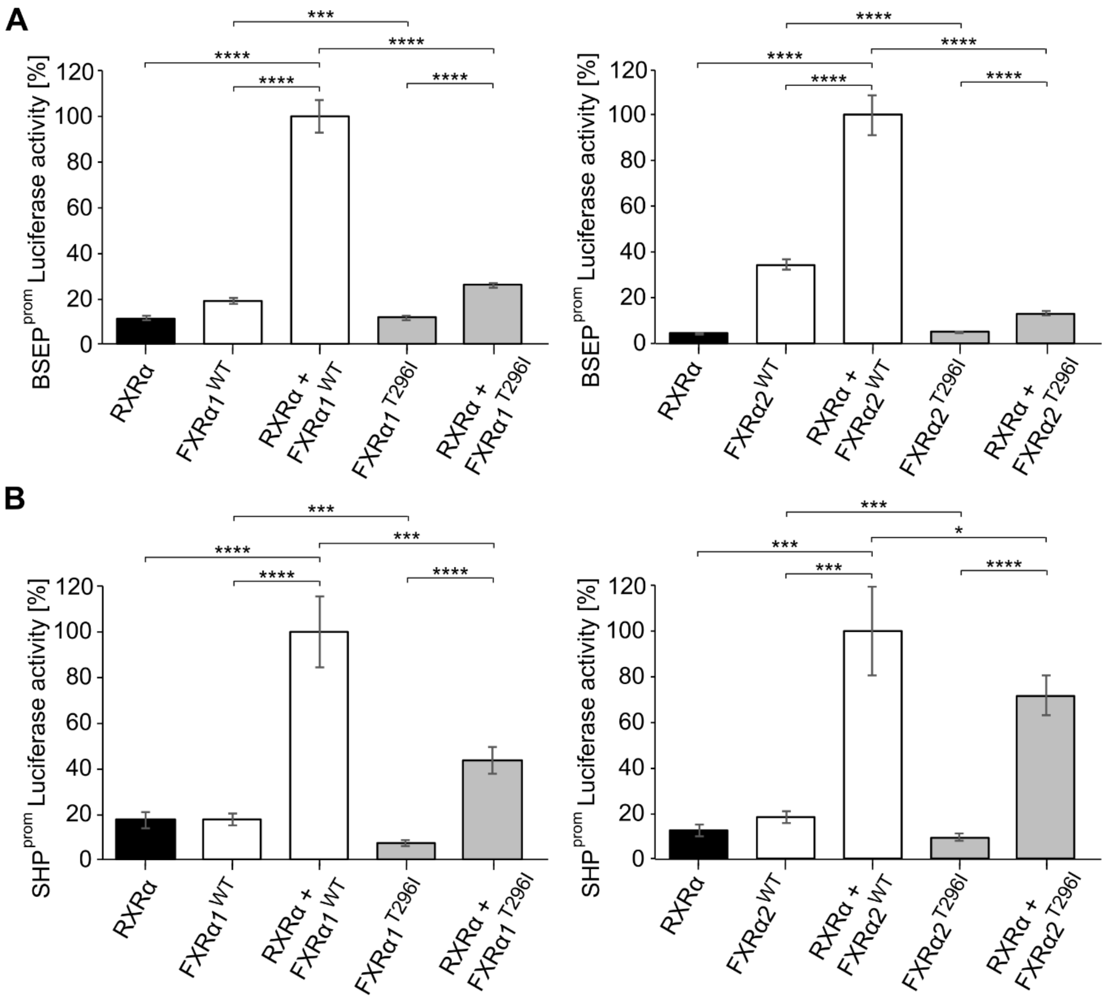


**Fig. S1: FXR T296I shows reduced transactivation of the BSEP and SHP promoters in transfected HEK293 cells.** A) Analysis of the luciferase enzymatic activity after transfection of HEK293 cells with a luciferase reporter gene containing the BSEP promoter ($BSEP^{prom}$-Luciferase) as well as different combinations of RXRα and either FXRα1$^{WT}$, FXRα2$^{WT}$, FXRα1$^{T296I}$, or FXRα2$^{T296I}$. The plasmid pRL-TK was included in each transfection for normalization. Cells were stimulated with an FXR and RXR ligand (OCA, 10µM and 9-cis-RA, 1µM). Each condition was assayed at least in three independent replicates. Values on the y-axis represent the mean and SD, expressed as % luciferase activity. The asterisks indicate a significant difference analyzed by a two-tailed Student t-test. B) Analysis of the luciferase enzymatic activity after transfection of HEK293 cells with a luciferase reporter gene downstream of the SHP promoter ($SHP^{prom}$-Luciferase) as well as different combinations of RXRα and either FXRα1$^{WT}$, FXRα2$^{WT}$, FXRα1$^{T296I}$, or FXRα2$^{T296I}$, as indicated on the x-axis. Further conditions and treatments are the same as described for panel A.

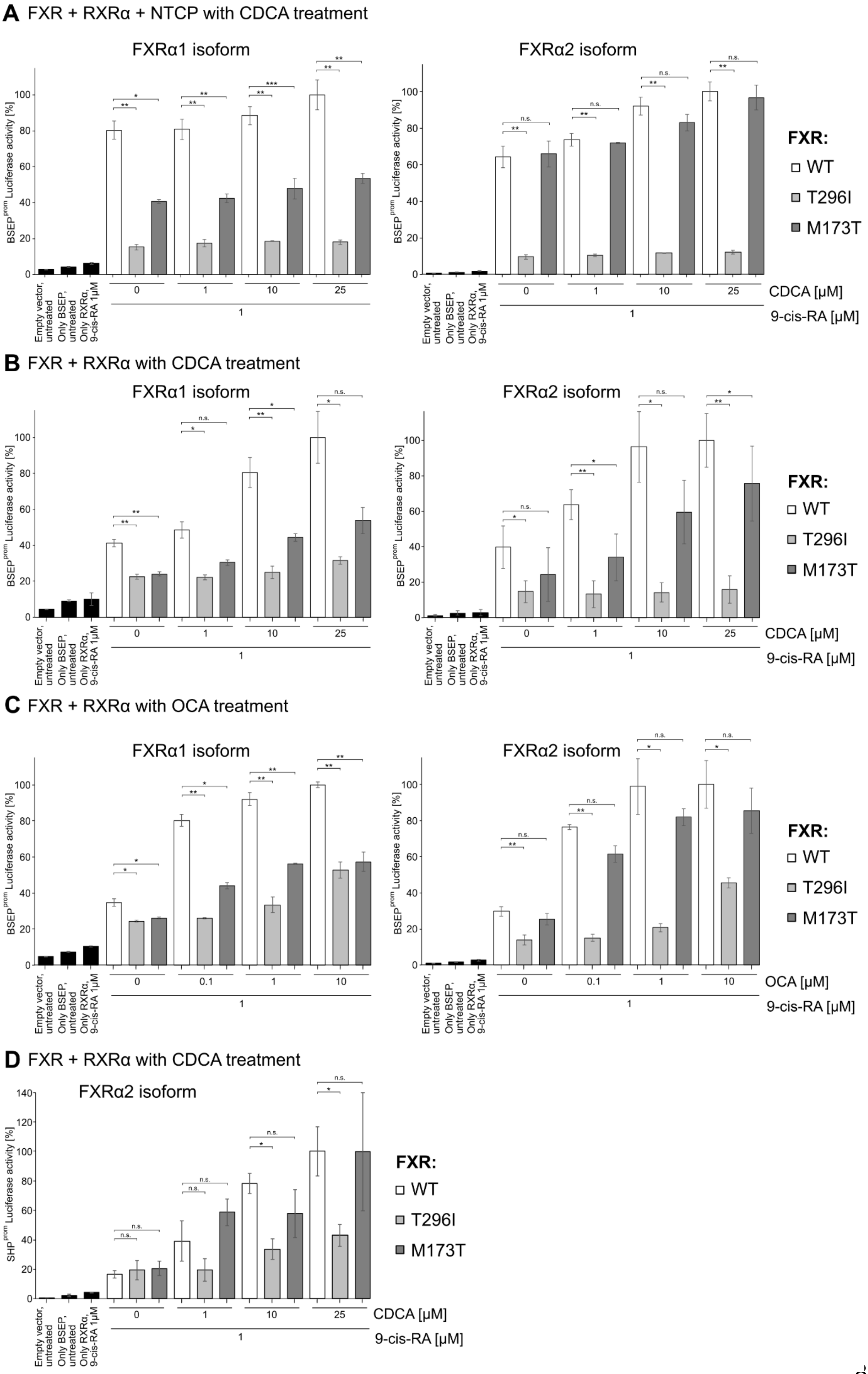
A FXR + RXRα + NTCP with CDCA treatment
FXRα1 isoform
FXRα2 isoform
BSEP$^{prom}$ Luciferase activity [%]
FXR:
WT
T296I
M173T
Empty vector, untreated
Only BSEP, untreated
Only RXRα, 9-cis-RA 1μM
CDCA [μM]
9-cis-RA [μM]
B FXR + RXRα with CDCA treatment
FXRα1 isoform
FXRα2 isoform
C FXR + RXRα with OCA treatment
FXRα1 isoform
FXRα2 isoform
OCA [μM]
D FXR + RXRα with CDCA treatment
FXRα2 isoform
SHP$^{prom}$ Luciferase activity [%]

**Fig. S2: FXR T296I shows reduced transactivation of the BSEP promotor in response to CDCA or OCA.** A) Analysis of the luciferase enzymatic activity after transfection of HEK293 cells with a luciferase reporter gene containing the BSEP promoter (BSEP$^{prom}$-Luciferase) as well as different combinations of RXRα and either FXRα1$^{WT}$, FXRα2$^{WT}$, FXRα1$^{T296I}$, FXRα2$^{T296I}$, FXRα1$^{M173T}$, or FXRα2$^{M173T}$. Additionally, cells were transfected with NTCP. Cells were stimulated with the RXR ligand (9-cis-RA, 1µM) and different concentrations of the FXR ligand CDCA (0, 1, 10, or 25µM). B) Analysis of the luciferase enzymatic activity after co-transfection using the BSEP$^{prom}$-Luciferase reporter system, RXRα, and either FXRα1$^{WT}$, FXRα2$^{WT}$, FXRα1$^{T296I}$, FXRα2$^{T296I}$, FXRα1$^{M173T}$, or FXRα2$^{M173T}$. Cells were stimulated with the RXR ligand (9-cis-RA, 1µM) and different concentrations of the FXR ligand CDCA (0, 1, 10, or 25µM). The data shown in the main text in Fig. 2, panel C and D highlights that part of the data that focusses on FXR WT and the T296I variant. C) Analysis of the luciferase enzymatic activity after co-transfection using the BSEP$^{prom}$-Luciferase reporter system, RXRα, and either FXRα1$^{WT}$, FXRα2$^{WT}$, FXRα1$^{T296I}$, FXRα2$^{T296I}$, FXRα1$^{M173T}$, or FXRα2$^{M173T}$. Cells were stimulated with the RXR ligand (9-cis-RA, 1µM) and different concentrations of the FXR ligand OCA (0, 0.1, 1, or 10µM). D) Analysis of the luciferase enzymatic activity after co-transfection using the FXR-regulated gene target SHP (SHP$^{prom}$-Luciferase) as reporter system, RXRα, and either FXRα2$^{WT}$, FXRα2$^{T296I}$, or FXRα2$^{M173T}$. Cells were stimulated with the RXR ligand (9-cis-RA, 1µM) and different concentrations of the FXR ligand CDCA (0, 1, 10, or 25µM). For all the experiments, values were obtained from three independent experiments. Values on the y-axis represent the mean and SD, expressed as % luciferase activity. For each experiment, the condition showing maximal luciferase activity was set to 100%. Statistical significance was analyzed by a two-tailed Student t-test.

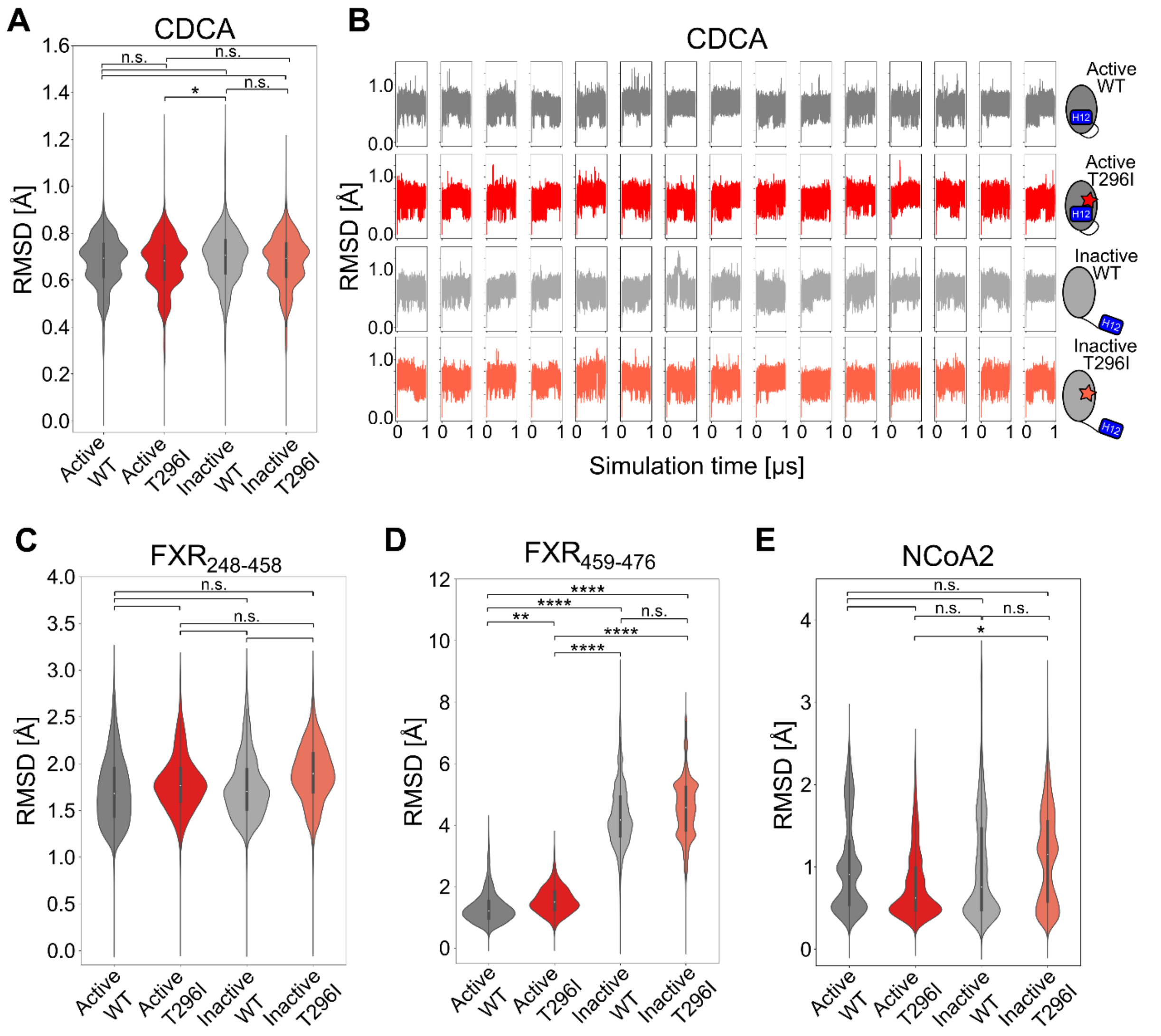


**Fig. S3: RMSD over MD simulation time of ligand, protein, and co-activation peptide.** (A) RMSD analysis of CDCA ligand (all atoms) over the MD simulation time (pooled over 15 replicas). (B) RMSD of CDCA ligand (all atoms) over the MD simulation time, shown for each of the 15 replicas. (C) RMSD of FXR LBD (residues 248-458) ($C_\alpha$ atoms), excluding the loop between helix (H) 11 and H12 and H12 itself (pooled over 15 replicas). (D) RMSD of FXR (residues 459-476) ($C_\alpha$ atoms), corresponding to the loop between helix 11 and helix 12 and helix 12 itself, over the MD simulation time (pooled over 15 replicas). (E) RMSD of NCoA2 co-activation peptide ($C_\alpha$ atoms) over the MD simulation time (pooled over 15 replicas). Significance tests were performed based on the means of the 15 replicas, respectively, using the Mann-Whitney U test. Violin plots were plotted using the Seaborn library[9], with the 1st to 3rd quartile within the box and the median marked as a white dot.

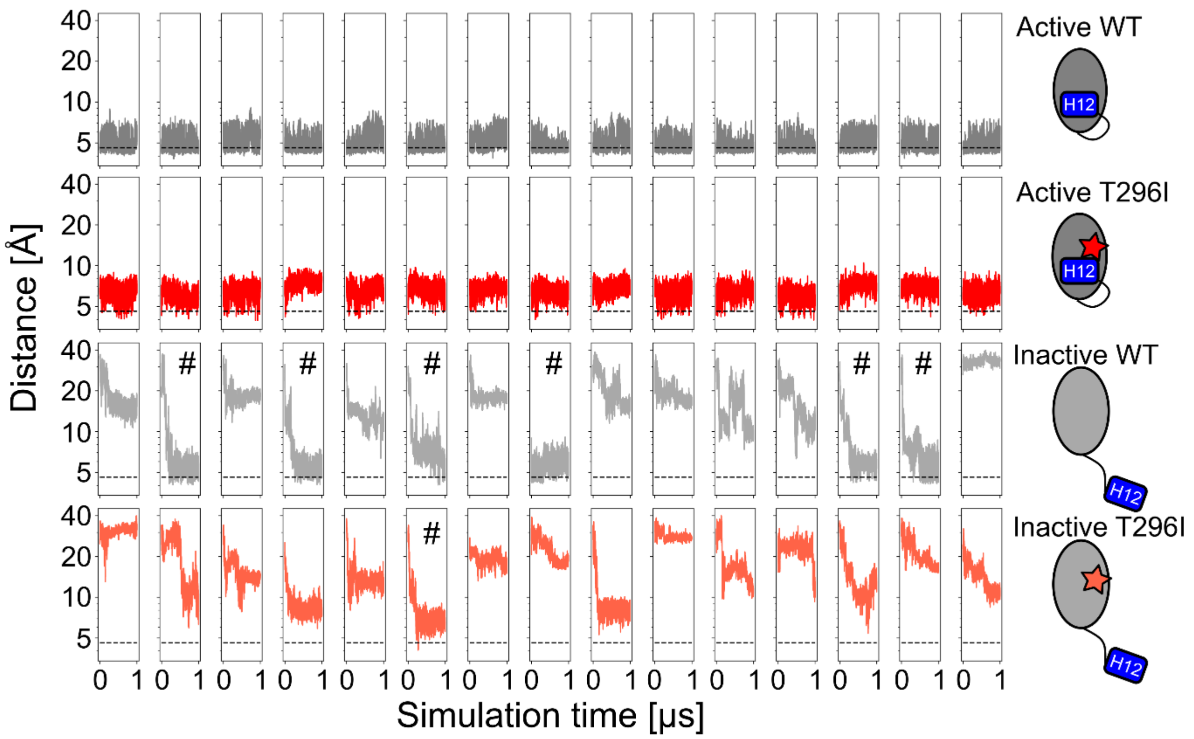


**Fig. S4: Distance between $C_{\beta}$ atoms of residues 296 and T466 over the MD simulations time.** The distances over the 15 independent replicas for the four different systems of FXR LBD are shown. The reference distance cutoff of 4.6Å, based on the crystal structure of the agonist-bound FXR LBD[10], is shown as a dashed line. For the inactive WT and inactive T296I systems, replicas are marked (#) where the reference distance cutoff was reached. Histograms and frequencies of occurrence shown in Fig. 3 were calculated based on this data.

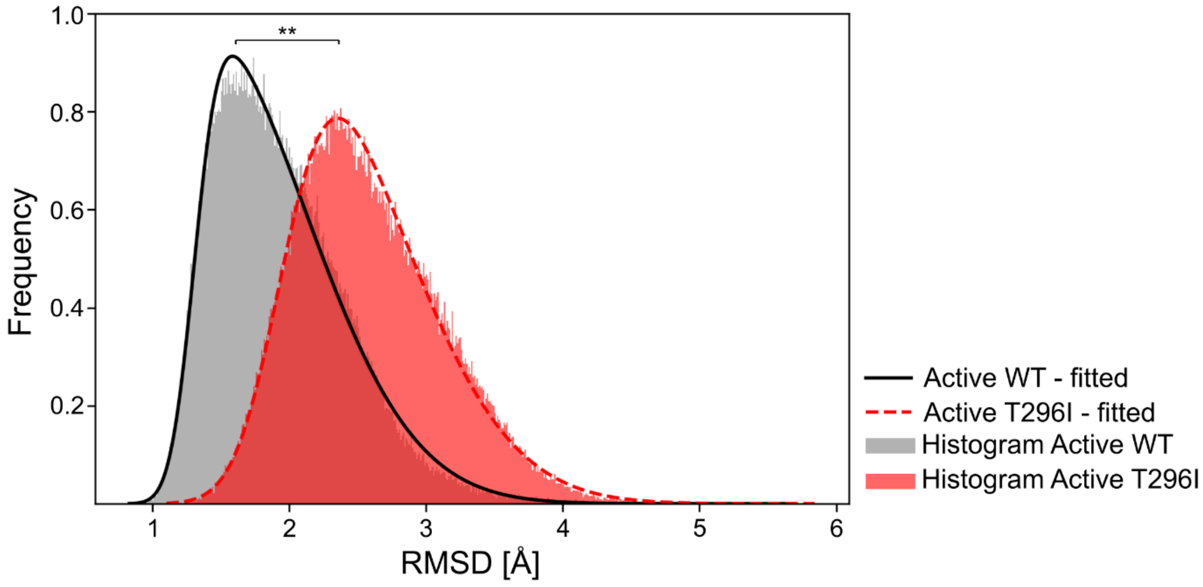


**Fig. S5: Skewed Gaussian fits on histograms of RMSD values of H12.** RMSD values of H12 and preceding T466 residue over 15 replicas of MD simulations with respect to the crystal structure of the agonist-bound FXR LBD are shown for active WT (grey) and active T296I (red). Using the scipy.stats module[11], skewed Gaussian functions were fitted (black and red lines). The fitted distributions are significantly different (assessed by a two-sided *t*-test based on the mean (active WT: 1.9Å, active T296I: 2.6Å) and the standard deviation (active WT: 0.5Å, active T296I: 0.5Å). The mean of the active WT was further used as a reference cutoff value for the expected RMSD fluctuations of H12 to the initial active structure. **: *p*-value = 0.0022.

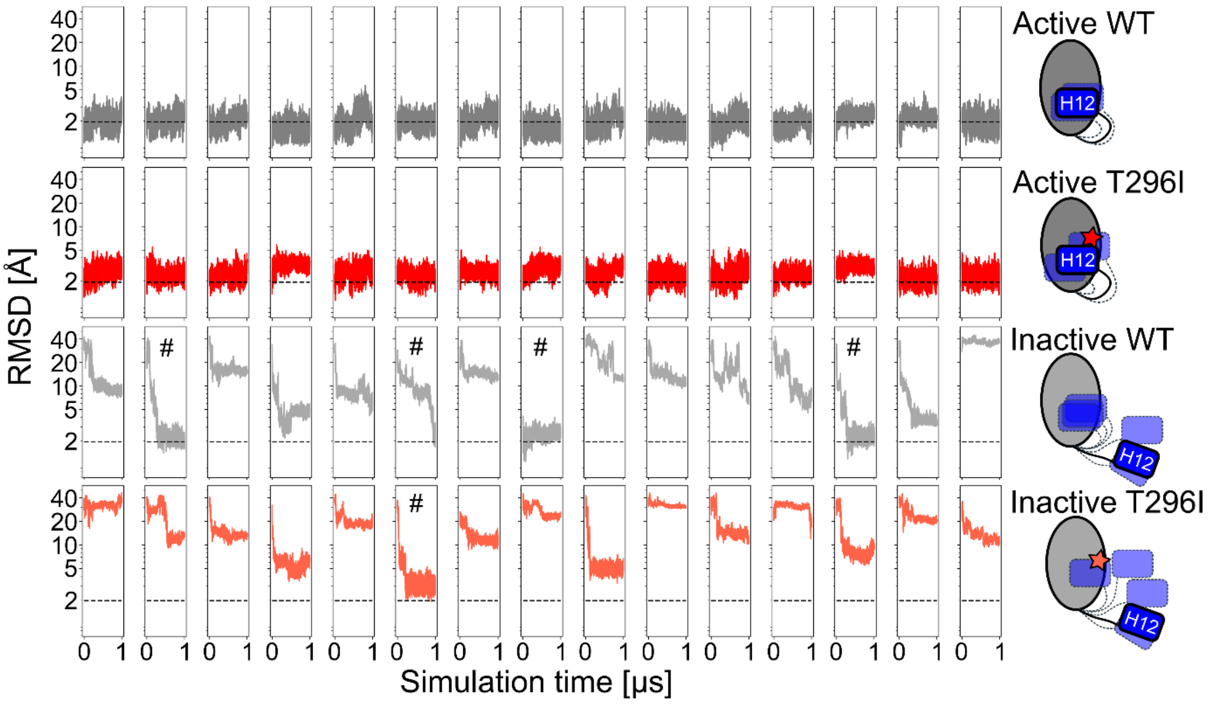


**Fig. S6: RMSD values of H12 over the MD simulation time.** The initial active WT structure, based on the crystal structure of agonist-bound FXR LBD[10], was used as a reference state for the RMSD analysis. The RMSD over the 15 independent replicas for the four different systems of FXR LBD is shown. The reference RMSD cutoff of 1.9Å, based on the mean of the fitted skewed Gaussian function for active WT values (Fig. S5), is shown as dashed lines. For the inactive WT and inactive T296I systems, replicas are marked (#) where the reference RMSD cutoff was reached. Histograms and frequencies of occurrence shown in Fig. 4 were calculated based on this data.

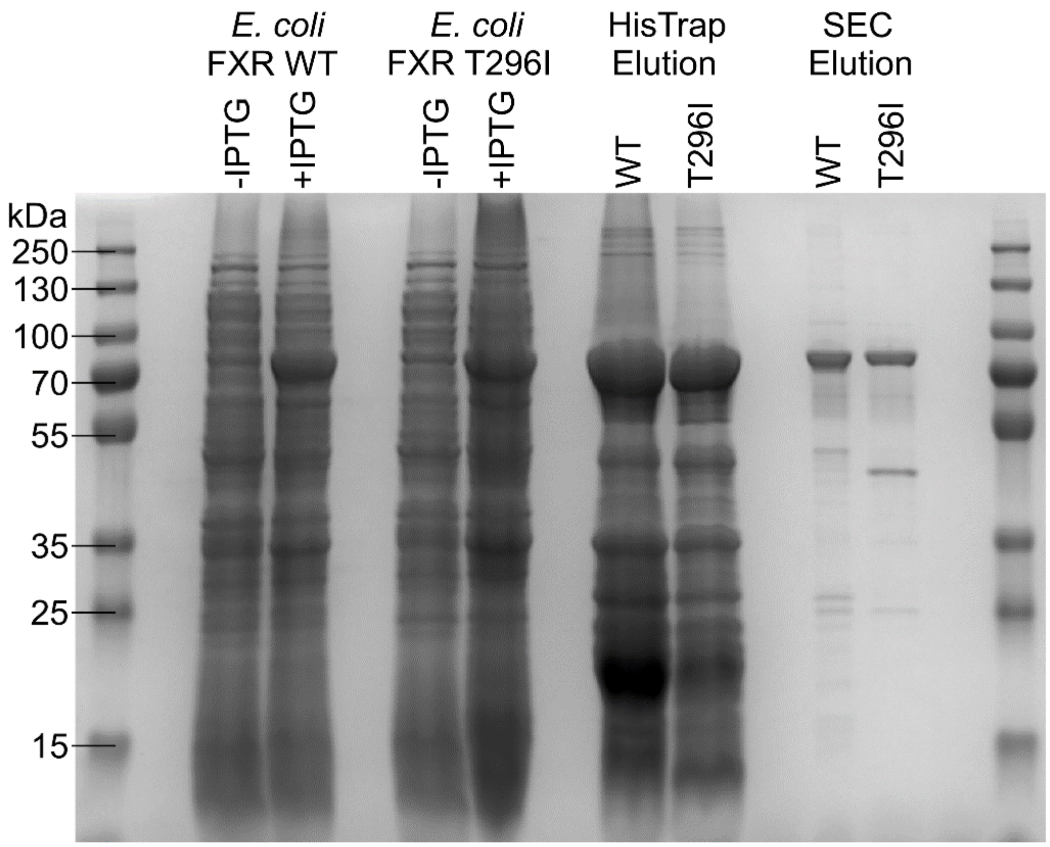


**Fig. S7: Coomassie staining of FXR WT and FXR T296I protein expression and purification.** The molecular weight of FXR protein with the SUMO and His-tag is ~70kDa. SEC: Size exclusion chromatography.

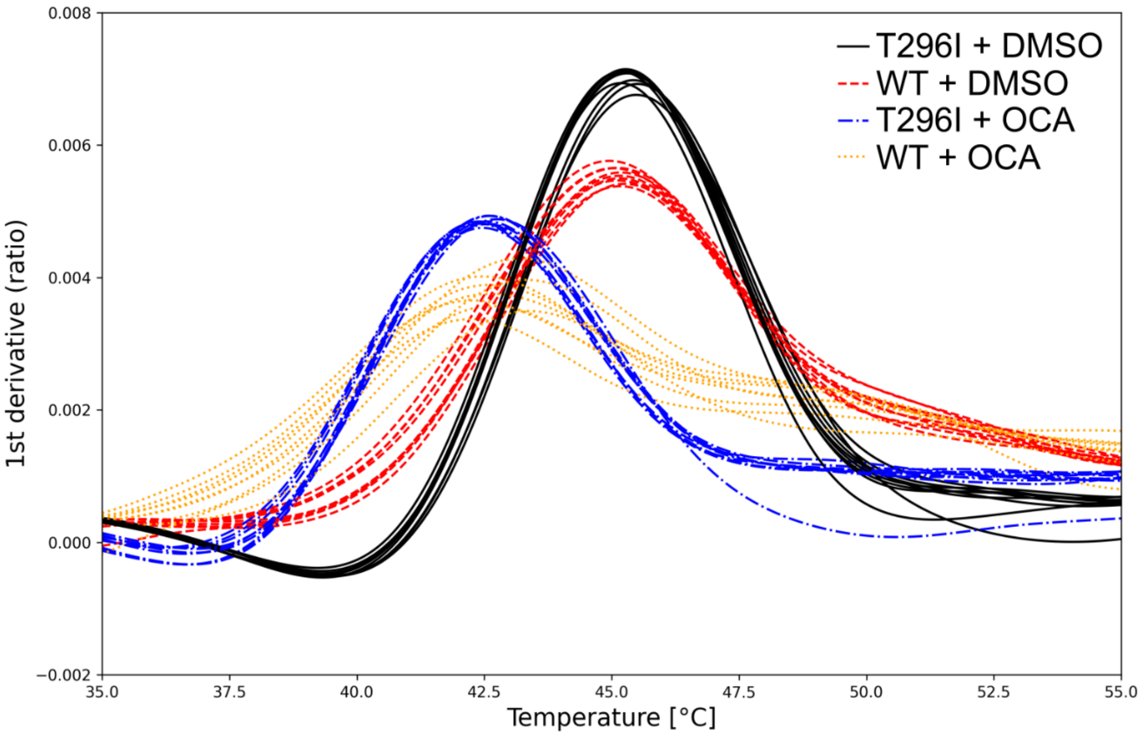


**Fig. S8: Thermostability assay of purified FXR WT and FXR T296I revealing binding of agonist OCA.** First derivative of fluorescence ratio 350nm over 330nm measured via nanoDSF. Measured melting temperatures and significance testing are shown Fig. S9.

| Significance | | Mean [°C] | Standard deviation [°C] |
|---|---|---|---|
| * / n.s. | FXR WT + DMSO | 45.17 | 0.09 |
| *** | FXR T296I + DMSO | 45.36 | 0.12 |
| n.s. | FXR WT + OCA | 42.84 | 0.41 |
| | FXR T296I + OCA | 42.58 | 0.12 |

**Fig. S9: Melting point measurements of FXR WT and FXR T296I.** Tm was measured via nanoDSF (Fig. S8) and statistical significance was tested using Welch's *t*-test. Protein samples were measured either in the absence (DMSO) or presence of agonist OCA.

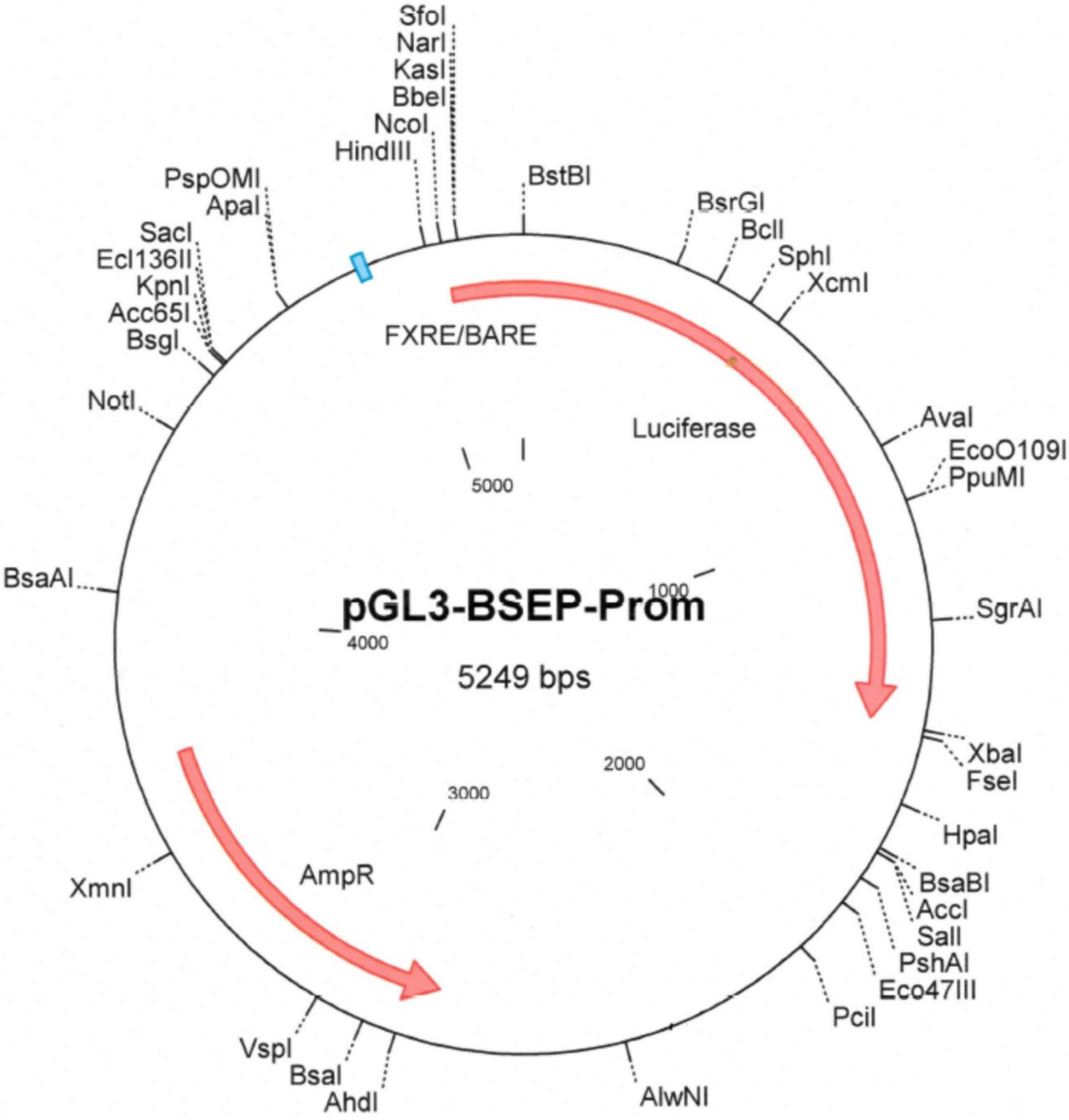


**Fig. S10: Plasmid map of the BSEP Promoter Luc construct.** Overview of the pGL3-BSEP-Promoter construct. Detailed sequences are available upon request.

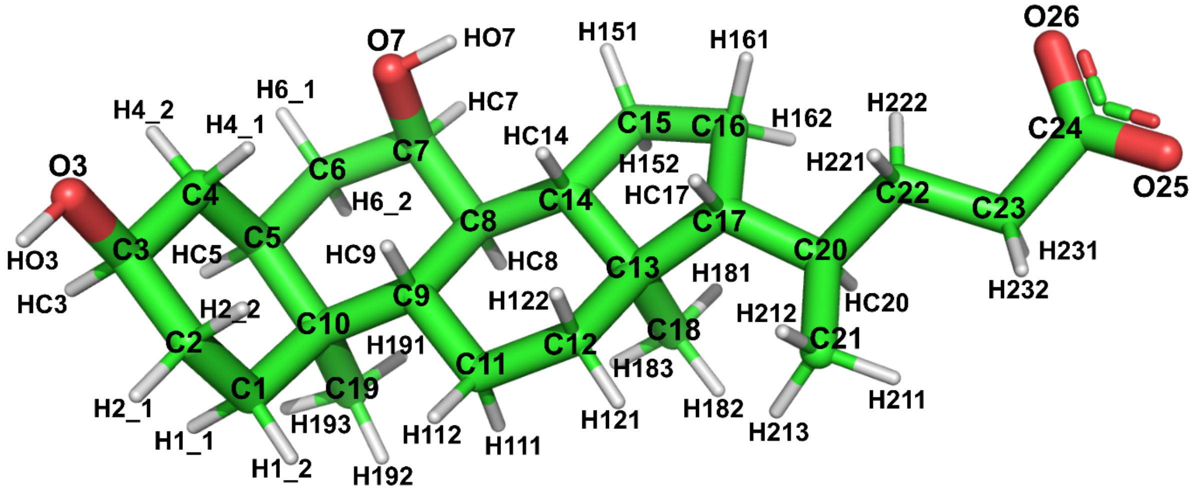


| Atom | Charge | Atom | Charge | Atom | Charge | Atom | Charge |
|---|---|---|---|---|---|---|---|
| C1 | -0.0476 | C7 | 0.1181 | C15 | -0.1328 | C21 | -0.0746 |
| H1_1 | 0.0044 | HC7 | 0.0410 | H151 | 0.0228 | H211 | 0.0044 |
| H1_2 | 0.0044 | O7 | -0.6947 | H152 | 0.0228 | H212 | 0.0044 |
| C2 | -0.0483 | HO7 | 0.4468 | C16 | -0.0703 | H213 | 0.0044 |
| H2_1 | 0.0171 | C8 | -0.0346 | H161 | 0.0529 | C22 | 0.0418 |
| H2_2 | 0.0171 | HC8 | 0.0557 | H162 | 0.0529 | H221 | -0.0266 |
| C3 | 0.1878 | C9 | 0.0268 | C17 | 0.0099 | H222 | -0.0266 |
| HC3 | 0.0129 | HC9 | 0.0527 | HC17 | 0.0100 | C23 | -0.0168 |
| O3 | -0.7098 | C10 | 0.0880 | C18 | -0.1016 | H231 | -0.0319 |
| HO3 | 0.4297 | C11 | -0.0192 | H181 | 0.0165 | H232 | -0.0319 |
| C4 | -0.0620 | H111 | 0.0115 | H182 | 0.0165 | C24 | 0.7860 |
| H4_1 | 0.0852 | H112 | 0.0115 | H183 | 0.0165 | O25 | -0.8101 |
| H4_2 | 0.0852 | C12 | -0.0520 | C19 | -0.0961 | O26 | -0.8101 |
| C5 | -0.0533 | H121 | -0.0008 | H191 | 0.0107 | | |
| HC5 | 0.0144 | H122 | -0.0008 | H192 | 0.0107 | | |
| C6 | -0.0347 | C13 | 0.0863 | H193 | 0.0107 | | |
| H6_1 | 0.0210 | C14 | 0.0017 | C20 | 0.0470 | | |
| H6_2 | 0.0210 | HC14 | 0.0098 | HC20 | -0.0038 | | |

**Fig. S11: Atomic point charges of the CDCA ligand used for MD simulations.** Top: Visualization of ligand structure and atom names. Bottom: RESP-derived atomic point charges.

### Supplementary Tables

**Table S1: Frequencies of occurrence of the distance between residue T296 and T466 ($C_\beta$ atoms) below the cutoff of 4.6Å, calculated for each MD simulations replica.**

| Replica # | Active WT[a] | Active T296I[a] | Inactive WT[a] | Inactive T296I[a] |
|---|---|---|---|---|
| 1 | 25.52 | 0.72 | 0.00 | 0.00 |
| 2 | 28.41 | 0.27 | 4.17 | 0.00 |
| 3 | 16.60 | 1.01 | 0.00 | 0.00 |
| 4 | 35.54 | 0.05 | 2.87 | 0.00 |
| 5 | 25.91 | 0.67 | 0.00 | 0.00 |
| 6 | 32.81 | 0.04 | 2.27 | 0.40 |
| 7 | 17.34 | 0.01 | 0.00 | 0.00 |
| 8 | 31.91 | 0.18 | 8.82 | 0.00 |
| 9 | 25.83 | 0.00 | 0.00 | 0.00 |
| 10 | 36.84 | 1.39 | 0.00 | 0.00 |
| 11 | 35.10 | 0.25 | 0.00 | 0.00 |
| 12 | 27.08 | 1.04 | 0.00 | 0.00 |
| 13 | 9.13 | 0.00 | 1.57 | 0.00 |
| 14 | 20.44 | 0.05 | 7.13 | 0.00 |
| 15 | 35.84 | 0.31 | 0.00 | 0.00 |
| Mean | 26.95 | 0.40 | 1.79 | 0.03 |
| STD | 8.20 | 0.45 | 2.85 | 0.10 |

[a] In %.

**Table S2: Frequencies of occurrence of the RMSD of H12 with respect to the initial active reference structure below the cutoff of 1.9Å, calculated for each MD simulations replica.**

| Replica # | Active WT[a] | Active T296I[a] | Inactive WT[a] | Inactive T296I[a] |
|---|---|---|---|---|
| 1 | 52.92 | 5.97 | 0.00 | 0.00 |
| 2 | 51.83 | 9.88 | 15.40 | 0.00 |
| 3 | 54.43 | 12.72 | 0.00 | 0.00 |
| 4 | 88.11 | 0.46 | 0.00 | 0.00 |
| 5 | 49.94 | 9.75 | 0.00 | 0.00 |
| 6 | 60.23 | 14.64 | 0.26 | 0.01 |
| 7 | 42.79 | 1.39 | 0.00 | 0.00 |
| 8 | 85.92 | 1.84 | 10.17 | 0.00 |
| 9 | 53.63 | 12.91 | 0.00 | 0.00 |
| 10 | 79.32 | 14.81 | 0.00 | 0.00 |
| 11 | 59.12 | 19.89 | 0.00 | 0.00 |
| 12 | 66.80 | 5.80 | 0.00 | 0.00 |
| 13 | 8.33 | 0.02 | 14.29 | 0.00 |
| 14 | 22.62 | 27.15 | 0.00 | 0.00 |
| 15 | 59.88 | 17.60 | 0.00 | 0.00 |
| Mean | 55.72 | 10.32 | 2.67 | 0.00 |
| STD | 21.15 | 7.90 | 5.59 | 0.00 |

[a] In %.

**Table S3: List of TaqMan™ Gene Expression Assays used for pre-amplification qPCR analysis.**

| Accession number | Gene Symbol | Taq ID | Amplicon length (base pairs) | Exon boundary | RefSeq |
|---|---|---|---|---|---|
| Hs01026590_m1 | NR1H4 | FXR1 | 78 | 10-11 | NM_001206979.1 |
| Hs01026592_m1 | NR1H4 | FXR2 | 84 | 2-3 | NM_001206979.1 |
| Hs00994824_m1 | ABCB11 | BSEP1 | 93 | 3-4 | NM_003742.2 |
| Hs00994811_m1 | ABCB11 | BSEP2 | 77 | 16-17 | NM_003742.2 |
| Hs00222677_m1 | NR0B2 | SHP | 87 | 1-2 | NM_021969.2 |
| Hs00983957_m1 | ABCB4 | MDR3 | 62 | 6-17 | NM_000443.3 |
| Hs00940446_m1 | RARA | NR1B1 | 68 | 6-7 | NM_000964.3 |
| Hs99999909_m1 | HPRT1 | HPRT1 | 100 | 6-7 | NM_000194.2 |
| Hs00188166_m1 | SDHA | SDHA | 70 | 5-6 | NM_001294332.1 |
| Hs02786624_g1 | GAPDH | GAPDH | 156 | 7 | NM_001256799.2 |

### Supplementary Movies

**Movie S1: Trajectory of inactive WT (replica no. 2), visualized over the entire MD simulations time.** The trajectory displays the conformational change from the inactive to the active state. For an easier judgment of the protein conformational state, the initial active state (based on the agonist-bound crystal structure) is depicted as green translucent representation. Important residues and motifs are highlighted (T296 as green sticks, T466 as cyan sticks, H12 in dark blue, NCoA2 peptide in magenta, and CDCA as pink sticks).

**Movie S2: Trajectory of inactive T296I (replica no. 6), visualized over the entire MD simulations time.** The trajectory displays the conformational change from an inactive to an almost active state. For an easier judgment of the protein conformational state, the initial active state (based on the agonist-bound crystal structure) is depicted as green translucent representation. Important residues and motifs are highlighted (I296 as orange sticks, T466 as cyan sticks, H12 in dark blue, NCoA2 peptide in magenta, and CDCA as pink sticks).

### Supplementary Methods

#### Plasmids, cloning and mutagenesis

The BSEP promoter plasmid based on pGL3-basic ($BSEP^{prom}$-Luc) (Fig. S10) was a kind gift from Roche. The human *SHP* promoter (bases -572 to +10, GenBank Accession Number AF044316)[12] was amplified by PCR using forward (5′-agggtaccTCCTAGACTGGACAGTGGGCAAAG-3′) and reverse (5′-gtgctagcCTTCCAGCTCTCTGGCTCTGTGTT-3′) to introduce, respectively, exogenous *KpnI* and *NheI* sites at 5´ends. Genomic DNA was extracted from human liver tissue, using the DNeasy® Blood & Tissue Kit (Qiagen) and was PCR amplified with the above primers. pCDNA3.1(+)-hRXR α[13] was obtained from Addgene (#135910). The FXR coding sequence was amplified from a human liver cDNA pool with the primer pair FXR-S1/-S2 (5´-ATGGGATCAAAAATGAATCTCATTGAACA-3´; 5´-TCACTGCACGTCCCAGATTTCACAGAG-3´) using Phusion HiFi DNA polymerase (ThermoFisher scientific). We thus obtained pCR2.1-FXR (α1 isoform, acc. no. NM_001206979.2). The FXR expression vector was constructed as follows: pmCherry-N1 (TaKaRa Bio) was linearized by PCR using primer pair pmCherry-tagdel-S1/-S2 (5´-CGGCCGCGACTCTAGATCATA-3´; 5´-GGTGGCGACCGGTGGATCCC-3´), removing the mCherry coding sequence in the process. FXR was amplified from pCR2.1-FXR using primer pair FXR-IFHD-S1/-S2 (5´-CCACCGGTCGCCACCATGGGATCAAAAATGAATCTCATTGAACA-3´; 5´-CTAGAGTCGCGGCCGTCACTGCACGTCCCAGATTTCACAGAG-3´), adding necessary terminal homologous overhangs. The α2 isoform (acc no. NM_005123.4) of FXR was generated from pnoCherry-FXR by inverse PCR using the primer pair FXR-MYTGΔ-S1/-S2 (5´-CTTGTTAACTGAAATTCAGTGTAAATCTAAGCGACTGAG-3´; 5´-CATTCAGCCAACATTCCCATCTCTTTGCATTTCC-3´) followed by phosphorylation of the 5´-termini and blunt-end self-ligation. The T296I variant was introduced into both FXR isoforms by site-directed mutagenesis using primer FXR-T296I-SDM (5´-TGACGGAAATGGCAATCAATCATGTACAGGTTCTT-3´) and the QuikChange Lightning Multi kit (Stratagene) according to the manufacturer's instructions. DNA sequencing was performed for all cDNAs used (Eurofins). Note that the numbering of the protein variant (T296I) is based on the alpha1 isoform (Uniprot acc. Q96RI1-1).

#### Luciferase Assay

Luciferase assays were performed using the Dual Luciferase reporter assay (Promega) according to the manufacturer's instructions. Briefly, HEK293 cells kept in DMEM containing

10% fetal calf serum (FCS) were seeded onto in 96-well plates at 7.000 cells per well and transfected the following morning with 100 ng of BSEP-Luc or SHP-Luc plasmid and 10 ng each of FXR and RXR expression plasmids using Fugene HD (Promega) at a 2.5:1 (reagent:DNA) ratio. Where indicated, cells were also transfected with 10ng of NTCP expression plasmid. Where applicable, plasmids were substituted with equal amounts of their respective empty backbones as control, and each well additionally received 5ng of pRL-TK as internal assay control (Fig.2, Fig.S2). Alternatively, cells were seeded onto 12-well plates at 150.000 cells per well and transfected the next morning with 1µg of the BSEP-Luc or SHP-Luc plasmid and 100ng each of FXR and RXR expression plasmids using Fugene HD (Promega) at a ratio of 2.5:1 (reagent:DNA). Where applicable, plasmids were substituted with equal amounts of their respective empty backbones as control, and each well additionally received 50ng of pRL-TK as internal assay control so that the total amount of DNA per well was always 1.25µg (Fig. S1). Each condition was assayed at least in three independent replicates. 4h after transfection, cells were pre-starved overnight by a medium change to DMEM containing 1% FCS (starvation medium) before stimulation with ligands in starvation medium for 24h. Cells transfected with FXRα1/2 expression plasmid were stimulated with CDCA or obeticholic acid (OCA, INT-747), cells transfected with RXRα were stimulated with 1µM 9-cis-retinoic acid (9-cis-RA), cells transfected with both FXRα1/2 and RXRα expression plasmids were stimulated with both ligands. Cells were washed with PBS, lysed in passive lysis buffer at RT for 20min and scraped into 1.5mL microcentrifuge tubes. Lysates were cleared by centrifugation at 16.000 g at 4°C for 10min, and supernatants were kept on ice. 10µL of each sample were assayed in duplicate using 50µL each of LARII and Stop & Glo reagents in a GloMax multi detection system (Promega). For the 96-well plates, cells were washed in PBS and lysed in 20 µl of passive lysis buffer at RT for 20 min. To facilitate cell lysis, 96-well plates were shaken at 500 rpm throughout the incubation period. Cells were then scraped with the tip of a 100 µl tip and 10 µl of each sample was transferred to a well of an opaque 96-well plate for the analysis. Each experimental condition was assayed in duplicate using 50 µL each of LARII and Stop & Glo reagents in a GloMax multi detection system (Promega).

**iPSC generation and hepatic differentiation**

Peripheral blood (one 7.5ml EDTA monovette) was obtained from the patient after informed consent and approval by the local ethics committee Hannover, Germany (2967/2015). The cells were layered on percoll and centrifuged 40min with 400xg at RT without brake and minimal acceleration. The interphase was removed and centrifuged. The pellet was resuspended in red cell lyses buff (0.15M $NH_4Cl$, 46mM $KHCO_3$, 0.5mM EDTA – all Roth) for 5min. Cells were washed and CD34+ hematopoietic stem cells were isolated using the CD34

MicroBead Kit (Miltenyi) according to manufacturer's protocol. Obtained cells were seeded in CD34+ media (StemSpan medium (Stemcell) supplemented with 2% Glutamine, 2% Penicillin/Streptomycin, 100ng/ml hSCF (Peprotech), 50ng/ml hTPO (Peprotech) and 100ng/ml hFlt3 (Peprotech). Cells were cultured for 24h and reprogrammed using commercial Sendai virus (CytoTune 2.0, Thermo Fisher Scientific) according to manufacturer's protocol. After 10-14 days, emerging iPSC-like colonies were picked from the initial plate and replated on CF1 feeder as fragments in 12-well cell culture dishes (TPP). Individual iPSC subclones were propagated by weekly picking single colonies and replated as fragments in 12-well plates for 6 weeks. At passage 6, clones were analyzed for Sendai virus clearance using a specific TaqMan assay according to manufacturer's protocol (Cytotune 2.0, Thermo Fisher Scientific) and only clones negative for Sendai virus underwent further characterization according to standard pluripotency assays. For the differentiation into hepatic organoids, iPSC colonies were split as single cells and seeded on GelTrex coated dishes (TPP) in E8 media containing no FGF2 and 10µM RHO-Kinase Inhibitor (Tocris) with a density of 120.000 cells/cm² to adapt for the subsequent differentiation protocol as described by Shinozawa et al.[14].

#### *In vitro* transcription and LNP production

FXRα2 and BSEP coding sequence was cloned in plasmid vector with modified T7 promoter, 5' and 3' UTR and segmented polyA tail. Plasmid was linearized by BspQI restriction digest. *In vitro* transcription was performed using HiScribe T7 mRNA kit with CleanCap Reagent AG (NEB) according to manufacturer's protocol and full substitution of UTP with me1Ψ-UTP (Jena Bioscience). Purification was performed using Monarch Spin RNA Cleanup Kit (NEB) according to manufacturer's protocol. LNP were produced by ethanol injection method. Briefly, lipids were dissolved in Ethanol at a molar ratio of 35:16:46:2.5 (C12 200, #LP-04-425, Corden Pharma; 1,2-dioleoyl-sn-glycero-3-phosphoethanolamine DOPE, #850725P Avanti; Cholesterol, #C8667, Sigma Aldrich; 1,2-Dimyristoyl-rac-glycero-3-methoxypolyethylenglycol-2000 DMG-PEG, #880151, Avanti) and total lipid concentration of 12.5 mM and mRNA was dissolved in 10 mM citrate buffer, pH4.0. LNPs were formulated by microfluidic mixing of the mRNA solution with the lipid mix at an N/P ratio of 6, a 3:1 flow rate ratio and 12 mL/min total flow rate in the NanoAssemblr® Ignite™ (Precision Nanosystems). Immediately after production, LNPs were dialyzed against PBS and concentrated using Amicon Ultra centrifugal filters with 10 kDa MWCO (UFC901024, Sigma-Aldrich). LNPs were supplemented with 5% (v/v) sucrose, frozen and stored at -80°C until transfection.

#### Functional assays of BSEP-mediated transport

At the end of the hepatic organoid differentiation, RHO-kinase inhibitor (Tocris) was added to the medium for an additional 1h at 37°C. Cell culture plates containing gel domes with organoids were placed on ice and supernatant was removed. Domes were washed twice with PBS and Cell Recovery Solution (Corning) was added in the middle of the domes. Plates were put on ice in a Styrofoam box that was mounted on an orbital shaker (40rpm) at 4°C. Within the next 45min, GelTreX should be dissolved. After GelTrex dissociation, the mixture was transferred to 14 ml ice-cold PBS and centrifuged for 10 min at 70 x g without brake. The obtained pellet was resuspended in ice-cold PBS and transferred into 1.5ml reaction tubes for assay or LNP transduction (500 ng total RNA per 1 million cells). For the Tauro-DBD assay, organoids were centrifuged 2min at 300 x g, supernatant was removed and 100µl media supplemented with 2µM of the fluorescent BSEP substrate Tauro-nor-THCA-24-DBD (Tauro-DBD) was added to each sample and incubated for given time points at 37°C. After incubation, samples were stopped with 1ml ice-cold PBS and centrifuged at 70 x g for 2 min, and for proper washing three cycles of supernatant removal and addition of 1ml ice-cold PBS were performed. The obtained organoids were finally seeded in 300µl 37°C pre-warmed HCM in a 24-well plate and analyzed using Olympus fluorescence microscope and CellSens software. Viewing fields (10x lens) containing 6-8 organoids were randomly selected in the phase contrast before fluorescence images for visualization of the luminal / canalicular accumulation of Tauro-DBD were acquired. ImageJ was used to calculate the ratio of the fluorescent area (fluorescence image) over the total organoid area (measured in phase contrast) from 7 viewing fields per condition.

**RNA preparation, reverse transcription, pre-amplification, and PCR analysis**

Total RNA was extracted from FFPE blocks of the patient's liver, one control liver, and two cirrhotic livers and purified using the AmoyDx FFPE DNA/RNA Kit (Amoy Diagnostics Co.) according to the manufacturer's instructions. The non-tumorous liver tissue from a patient undergoing liver metastasis resection was used as control tissue and histopathologically contained no signs of hepatitis or fibrosis (denoted as healthy control). Cirrhotic tissue was obtained from two patients undergoing resection of hepatocellular carcinoma. The cirrhotic tissue used was histopathologically free of HCC. The study was approved by the local ethics committee Magdeburg, Germany (33/01). Purified RNA was eluted with 30µL nuclease-free water. RNA integrity was assessed by microcapillary electrophoresis on 2100 BioAnalyser (Agilent Technologies). 100ng of RNA was reverse transcribed for 120min at 37°C followed by 5min of enzyme inactivation at 85°C using the High Capacity cDNA Reverse Transcription Kit (Applied Biosystems) with the following components: 1xRT buffer, 100mM dNTP mix, 1xRandom Primers, 50 U of MultiScribe Reverse Transcriptase and 1 U of RNAse inhibitor.

cDNA was then diluted 1:3 in nuclease-free water. Before qPCR, sequences of interest in the cDNA were pre-amplified through 10 cycles using 1xTaqMan PreAmp Master Mix (Applied Biosystems) and a pool of TaqMan Gene Expression Assays (Table S3), diluted 1:200 in nuclease-free water, in a final volume of 50µL (0.05x each Assay). Relative quantification of mRNA was performed according to the comparative $2^{-\Delta\Delta CT}$ method with SDHA as an endogenous control. HPRT1 and GAPDH were excluded as endogenous controls due to high Ct values. All TaqMan Gene Expression Assays used for cDNA pre-amplification and qPCR amplification were ordered from Applied Biosystems.

**Setup of MD simulations and production replicas**

The structural models were prepared for molecular dynamics (MD) simulations using the AMBER21 package[15]. Overall, four different systems were prepared: FXR wildtype in the active state (hereafter termed "active WT"), FXR T296I variant in the active state ("active T296I"), FXR wildtype in the inactive state ("inactive WT"), and FXR T296I variant in the inactive state ("inactive T296I"). Maestro (Schrödinger, LLC, New York) was used for assigning protonation states with PROPKA[16] at pH 7.0; histidine HIP states were reverted to HIE. Parameters for the protein were taken from the ff14SB force field[17], and the TIP3P[18] parameters were used for the water and ions. The protein was solvated in a cubic water box, and $Na^+$ ions were added to neutralize the protein charges using tleap[15]. The CDCA ligand was parametrized in its physiologically relevant deprotonated form. Electrostatic point charges were obtained with the RESP method[19] to represent the electrostatic potential of the ligand using the R.E.D. server (PyRED version April 2022)[19–22] with the electrostatic potential calculated at the 6-31G(d) level of theory using Gaussian16 vC.01 (Gaussian, Inc., Wallingford, USA). In line with the deprotonated state of the carboxylic acid group, the overall molecule charge was kept at -1. The ligand, as well as the derived point charges, are depicted in Fig. S11. Applying the SHAKE algorithm[23] to constrain bond lengths of hydrogen atoms to heavy atoms enabled a time step of 2 fs. Long-range electrostatic interactions were considered using the particle mesh Ewald algorithm[24]. Fifteen independent replicas were set up for each system, and each system was minimized for 1000 steps using the steepest descent algorithm, followed by 1000 steps using the conjugate gradient algorithm, applying harmonic positional restraints with a force constant of 50 kcal $mol^{-1}$ $Å^{-2}$ to the system excluding the protein hydrogens, water molecules, protein side chains, protein, and ligand atoms in consecutive runs. The system was heated to 300 K for 25 ps in the NVT ensemble using the Langevin thermostat[25] with a collision frequency of 2.0 $ps^{-1}$ with the protein atoms restrained with a force constant of 10 kcal $mol^{-1}$ $Å^{-2}$. Within the following seven consecutive equilibration steps performed in the NPT ensemble using the Berendsen barostat[26] at 1 bar, the restraints were removed (after 100 ps of the overall 4975 ps simulation time). Using hydrogen mass repartitioning[27], the time step was increased to 4 fs for the production replicas (each replica was simulated for 1 μs, giving a total length of 15 μs per system). Coordinates were stored in time steps of 100 ps.

#### Analysis of MD simulations

Postprocessing and analysis of the MD trajectories were performed with CPPTRAJ[28] implemented in AmberTools21[15]. Root mean square deviations (RMSD) of the systems over the production time were based on $C_{\alpha}$ atom positions (for FXR and NCoA2) or all atoms (for CDCA) to analyze the structural variability of the systems. Analysis of the RMSD of H12 and the preceding T466 (residues 466 to 473) was performed against the reference of the active state of the initial FXR WT structure, first fitting the conformations along a trajectory on the most stable core of the four different FXR systems (active WT, active T296I, inactive WT, inactive T296I) over the production time. In detail, frames were extracted from all trajectories every 10ns and analyzed using the BIO3D package[29–31] to identify the least mobile residues throughout the simulations, which resulted in a part of helix 4 (residues 328 to 335) located at the core of the FXR LBD. The distance between the $C_{\beta}$ atoms of residue 296 and T466 was measured to avoid a bias due to side-chain motions. Visualization was done using PyMOL v2.4.0. (Schrödinger, LLC, New York) or VMD v1.9.3.[32]. The distance between H12 (residues 467 to 473) and LBD core (residues 248 to 459) was measured using the center of mass function within PyMOL.

#### Statistical Analysis of MD simulations

Significance tests were performed using the Mann-Whitney U test if not indicated otherwise. The indicated significance levels are n.s. (not significant): $p > 0.05$, *: $p \leq 0.05$, **: $p \leq 0.01$, ***: $p \leq 0.001$, ****: $p \leq 0.0001$.

#### Bacterial expression plasmid construction

In order to express the FXRα2 isoform (residues 1 to 472, uniprot acc. Q96RI1-2) in *Escherichia coli* (*E. coli*), the FXRα2 gene was PCR-amplified (Q5 Polymerase, NEB, Ipswich, USA) from the pnoCherry::fxr_alpha2_wt plasmid with the following primers: FXR_fw 5'-GAACAGATTGGTGGTATGGGATCAAAAATGAATCTC-3' and FXR_rv 5'-CAGCCGGATCTCACTGCACGTCCCAGATTTC-3'. The pET SUMO vector backbone (Invitrogen, Waltham, USA) was PCR-amplified using the following primers: pET SUMO_fw 5'-TGAGATCCGGCTGCTAACAAAGCC-3' and pET SUMO_rv 5'-ACCACCAATCTGTTCTCTGTGAGC-3'. PCR products were run on agarose gels for expected size verification, DpnI (NEB) digested for 1h at 37°C and purified using NucleoSpin Gel and

PCR clean-up kit (Macherey-Nagel, Dueren, Germany). For homologous recombination, vector backbone and insert were used at a ratio of 1:3 for transformation of competent *E. coli* DH5alpha (DE3) (Invitrogen, Waltham, USA). Transformed cells were plated on Luria-Bertani agar plates containing 50µg/mL kanamycin (Sigma Aldrich, St. Louis, USA) and incubated overnight at 37°C. Plasmid isolation of single colonies was performed using the NucleoSpin Plasmid kit (Macherey-Nagel) according to manufacturer's instruction and sequenced (Eurofins, Luxembourg). Recombinant plasmid (pET SUMO::fxr_alpha2_wt) was further used for the transformation of competent *E. coli* Rosetta (DE3) pLysS cells (Sigma Aldrich) for protein expression. The T296I mutation was introduced via a modified QuikChange protocol taking the pET-SUMO-FXR_alpha2_wt as template, using the primers: FXR_T296I_fw 5'-GACGGAAATGGCAattAATCATGTACAGG-3' and FXR_T296I_rv 5'-CATGATTaatTGCCATTTCCGTCAAAATG-3' (small letters indicating codon exchange). Similar to the wildtype, variant recombinant plasmid pET-SUMO::fxr_alpha2_T296I was first subcloned in *E. coli* DH5alpha cells, sequenced to verify the site-specific exchange, and used for transformation of competent *E. coli* Rosetta (DE3) pLysS for protein expression.

**Protein expression and purification**

FXR WT and T296I protein expression was performed with *E. coli* Rosetta(DE3) pLysS pET-SUMO::fxr_alpha2_wt or pET-SUMO::fxr_alpha2_T296I grown in terrific broth (TB) medium with 50µg/mL kanamycin (Sigma Aldrich). Cultures were grown at 37°C and shaking at 180 rpm until reaching an OD600 of 0.9. Protein expression was induced with 1mM of IPTG (Sigma Aldrich, St. Louis, USA) and cultures were further incubated at 20°C for 20h at 180 rpm. Bacterial cells were harvested by centrifugation (9,150xg for 12min at 4°C (Avanti JXN-26, Beckman Coulter, Brea, USA)) and cell pellets either stored at -80°C or directly used for subsequent cell lysis. For cell lysis, the bacterial pellet was resuspended in 10mL/g Lysis Buffer (50mM $Na_2HPO_4$, 300mM NaCl, pH 8.0, supplemented with protease and phosphatase inhibitors (Complete tablets, Roche, Basel, Switzerland)), 1mg/mL of lysozyme (Sigma Aldrich) and 5µg/mL of DNAse 1 (Roche, Basel, Switzerland) were added. The mixture was then sonicated on ice three consecutive times for 15min each with an interval pulse of 2 seconds (UP200St Sonicator, Hielscher Ultrasonics, Teltow, Germany). Cell debris was removed by centrifugation at 58,540xg for 60min at 4°C (Avanti JXN-26, Beckman Coulter, Brea, USA) before loading the supernatant on a pre-equilibrated HisTrap column (HisTrap HP, Cytiva, Marlborough, USA). Pre-equilibration was performed with binding buffer (50mM $Na_2HPO_4$, 300mM NaCl, 30mM imidazole (Thermo Fisher Scientific, Waltham, USA), pH 8.0). After protein extract application, the column was washed with 4 column volumes of binding buffer, before eluting the recombinant protein with elution buffer (50mM $Na_2HPO_4$, 300mM NaCl,

500mM imidazole, pH 8.0). The purity of the recombinant protein was further improved via size exclusion chromatography (SEC) using a HiLoad 16/600 Superdex 200pg column (Cytiva, Marlborough, USA, product number 28989335) in SEC buffer (50mM $Na_2HPO_4$, 300mM NaCl, pH 8.0). Samples for SDS-PAGE analysis (12% gels, MiniProtean Gel, BioRad, Hercules, USA) were taken before and after induction, after HisTrap elution, and after SEC elution (Fig. S7). Fractions containing purified protein were concentrated and stored at -80°C in 50µM aliquots.

**Thermostability assay**

Recombinant FXR WT and FXR T296I protein samples were analyzed for their thermostability using the nanoDSF technology (Prometheus, Nanotemper, Munich, Germany). Protein samples were analyzed at a concentration of 25µM in SEC buffer (50mM $Na_2HPO_4$, 300mM NaCl, pH 8.0) either in the presence of 10-fold excess of the agonist OCA (INT-747, AbMole, Houston, USA) (250µM in 2.5% DMSO) or with a respective 2.5% DMSO control. Prometheus Standard capillaries (Nanotemper, Munich, Germany; with three technical replicates per sample) were used and three runs from 20°C to 90°C with an increase of 1°C per minute were performed (Fig. S8 and Fig. S9).
